# Functional Connectivity Differences in Distinct Dentato-Cortical Networks in Alzheimer’s Disease and Mild Cognitive Impairment

**DOI:** 10.1101/2024.02.02.578249

**Authors:** Ivan A. Herrejon, T. Bryan Jackson, Tracey H. Hicks, Jessica A. Bernard, Alzheimer’s Disease Neuroimaging Initiative

## Abstract

Recent research has implicated the cerebellum in Alzheimer’s disease (AD), and cerebro-cerebellar network connectivity is emerging as a possible contributor to symptom severity. The cerebellar dentate nucleus (DN) has parallel motor and non-motor sub-regions that project to motor and frontal regions of the cerebral cortex, respectively. These distinct dentato-cortical networks have been delineated in the non-human primate and human brain. Importantly, cerebellar regions prone to atrophy in AD are functionally connected to atrophied regions of the cerebral cortex, suggesting that dysfunction perhaps occurs at a network level. Investigating functional connectivity (FC) alterations of the DN is a crucial step in understanding the cerebellum in AD and in mild cognitive impairment (MCI). Inclusion of this latter group stands to provide insights into cerebellar contributions prior to diagnosis of AD. The present study investigated FC differences in dorsal (dDN) and ventral (vDN) DN networks in MCI and AD relative to cognitively normal participants (CN) and relationships between FC and behavior. Our results showed patterns indicating both higher and lower functional connectivity in both dDN and vDN in AD compared to CN. However, connectivity in the AD group was lower when compared to MCI. We argue that these findings suggest that the patterns of higher FC in AD may act as a compensatory mechanism. Additionally, we found associations between the individual networks and behavior. There were significant interactions between dDN connectivity and motor symptoms. However, both DN seeds were associated with cognitive task performance. Together, these results indicate that cerebellar DN networks are impacted in AD, and this may impact behavior. In concert with the growing body of literature implicating the cerebellum in AD, our work further underscores the importance of investigations of this region. We speculate that much like in psychiatric diseases such as schizophrenia, cerebellar dysfunction results in negative impacts on thought and the organization therein. Further, this is consistent with recent arguments that the cerebellum provides crucial scaffolding for cognitive function in aging. Together, our findings stand to inform future clinical work in the diagnosis and understanding of this disease.

## Introduction

Alzheimer’s disease (AD), the most common neurodegenerative disorder, disrupts the quality of life of older adults by impairing cognition^1,2^ and motor functions.^3–5^ In addition to groundbreaking work focused on the hippocampus and medial temporal lobes, researchers have historically focused on the cerebral cortex in attempts to understand these symptoms and the etiology of the disease. However, despite structural connectivity from the cerebellum to cortical regions prone to atrophy in AD^6,7^, the cerebellum was thought to be spared in AD, in part due to it being historically regarded as a motor structure.^2,8–10^

Lesion studies, clinical, and neuroimaging research have implicated the cerebellum in motor and non-motor functions including reward processing, executive function, learning, emotion, language, memory, somatosensory processing, and attention.^11–13^ Research on animal models of AD and patients with AD has shown deficits in these cognitive and motor processes and cortical atrophy in the areas associated with their modulation.^3,14^ Notably however, the cerebral cortex receives projections from the cerebellar dentate nucleus (DN) via the thalamus forming at least two distinct closed loops. The dorsal dentate nucleus (dDN) projects to motor regions (i.e. primary motor cortex) and is associated with motor functions, while the ventral dentate nucleus (vDN) projects to cognitive areas (i.e. prefrontal cortex (PFC)) and is associated with cognitive functions.^15–19^ Emerging evidence for a third region has been seen in resting state connectivity ^20^ though to this point no tractography work has provided information on cortical targets. Thus, for our purposes here we are focusing on the dorsal and ventral aspects of the dentate.

Recently, however, a growing literature highlights the potential importance of the cerebellum in AD. Pathology present in the hippocampus or cortical regions^21,22^ including gray and white matter atrophy, amyloid plaques, and tau, as well as functional connectivity (FC) differences have also been found in the cerebellum. For example, cerebellar gray (GM) and white matter (WM) are negatively impacted in AD.^5,23–27^ More recently, Gellersen and colleagues^10^ conducted a meta-analysis investigating cerebellar GM atrophy in cognitively normal older adults (CN) and AD and mapped it onto the cerebellum using three approaches: comparing it against network connectivity patterns, functional boundaries based on a battery of 26 tasks^12^, and along major functional gradients. In AD, like in CN, there was GM loss associated with the frontoparietal and the default mode networks (DMN). In the gradient analyses, cerebellar structures associated with cognitive functions experienced atrophy, but not motor ones. With respect to territories associated with task-based functional activation, in AD, GM loss was present in working memory, attention, and language processing areas of the cerebellum. Interestingly, there was minimal overlap between the areas that experienced GM loss in CN and AD, but in both groups, there was a bias towards cerebellar atrophy in cerebellar regions associated with cognition. Finally, in AD, cerebellar GM atrophy was predominantly present in the right hemisphere only.^10^

In work using resting state functional connectivity (FC), Olivito and colleagues^26^ demonstrated cerebellar FC differences in AD compared to HC.^26^ Using seeds in the left and right DN, they found higher FC between the DN and the lateral temporal cortex in AD compared to CN individuals. As expected, AD patients had worse outcomes in memory scores, though performance was not related to FC differences. Finally, this higher connectivity in AD was postulated as a possible compensatory mechanism. However, Olivito investigated the dentate nucleus as a whole, despite evidence suggesting these regions are part of distinct networks.^15,17,18^ As such, whether the dDN and vDN networks differ in AD, and the degree to which they may relate to cognitive and motor function, are open questions.

In this context of growing interest in understanding the cerebellum in AD, the present study investigated FC differences in dDN and vDN networks to characterize the effects of AD on these cerebellar networks. Additionally, we included MCI participants to understand FC differences with cognitive decline prior to the onset of a formal diagnosis of disease. Finally, we included cognitive and motor task measures to test whether FC was associated with task performance. We hypothesized that (i) due to the pathological impact of AD on the volume of functionally-connected cerebellar and cerebral regions and the noted changes in cerebrocerebellar functional connectivity in MCI and AD^2,5,26,28^, we will find increased functional connectivity within the dorsal and ventral dentato-cortical networks in MCI compared to CN and a decrease in AD participants compared to MCI; (ii) task performance will track with severity of cognitive decline and both motor and cognitive scores will be lower in AD than in those with MCI, which will, in turn, be lower than CN; (iii) task performance associations with FC will follow network dysfunction and thus functional connectivity of the dDN network will be associated with performance on motor tasks and the vDN network will be associated with performance on cognitive tasks in both MCI and AD groups.

## Methods

The current study and planned analysis were pre-registered on the Open Science Framework prior to initiation (OSF; https://doi.org/10.17605/OSF.IO/N5DM9). Data used in the preparation of this manuscript were obtained from the Alzheimer’s Disease Neuroimaging Initiative (ADNI) database (adni.loni.usc.edu). ADNI was launched in 2003 as a public-private partnership, led by Principal Investigator Michael W. Weiner, MD. The primary goal of ADNI has been to test whether serial magnetic resonance imaging (MRI), positron emission tomography (PET), other biological markers, and clinical and neuropsychological assessment can be combined to measure the progression of MCI and early AD.

### Participants

Five hundred forty-seven participants were selected from the ADNI 3 dataset. We used the LONI advanced search tool to select all ADNI 3 participants with a complete set of images (at least one complete resting-state bold EPI, field maps, a high-resolution T1-weighted structural image (MPRAGE), a T2*-weighted, and a FLAIR) and all cognitive task scores. Raw scores obtained via the Wechsler Memory Scale-III Logical Memory task, which requires participants to repeat a story immediately after hearing it and the number of correctly generated story items resulted in the immediate recall score; the process of repeating the story again after a 20-30 minute delay with the same scoring technique resulted in the delayed recall score. For further task descriptions and details see: https://adni.loni.usc.edu/wp-content/themes/freshnews-dev-v2/documents/clinical/ADNI3_Protocol.pdf. Behavioral motor measures were scores from neurological exams investigating motor processes such as gait and cerebellar finger to nose performance (for task descriptions and details see: https://adni.loni.usc.edu/wp-content/uploads/2012/10/ADNI3-Procedures-Manual_v3.0_20170627.pdf). An initial filter in LONI was conducted to include subjects that had scan, motor, and cognitive data, leaving four hundred seventy-eight participants with the following breakdown based on group: AD, N=47; MCI, N=139; and CN, N=292. For more information regarding the sample please refer to Table 1. Quality control information provided by ADNI was used. Resting state scans were an average of 7-12 minutes long. Additional information regarding acquisition parameters can be found in https://adni.loni.usc.edu/methods/mri-tool/mri-analysis.

**Table 1.** Demographic, cognitive, motor, and motion parameters Means (Standard Deviation) information of CN, MCI, and AD groups. Statistical comparisons between groups are reported in text.

|  | CN | MCI | AD |
| --- | --- | --- | --- |
| Age (years) | 72.2(7.31) | 74.7(8.21) | 76.7(8.52) |
| Age Range (years) | 55-95 | 55-97 | 55-95 |
| Education | 16.8(2.2) | 15.8(2.7) | 15.1(2.4) |
| Sex (M/F) | 118(174) | 74(65) | 25(22) |
| Sex (%Female) | 59.70% | 46.40% | 44.70% |
| African American (%) | 8.87% | 7.14% | 6.38% |
| Asian (%) | 2.73% | 0.71% | 2.14% |
| Caucasian (%) | 84.64% | 90.00% | 82.98% |
| Multiracial (%) | 2.73% | 0.71% | 6.38% |
| Native American (%) | 0.34% | 0.00% | 0.00% |
| Unknown = (%) | 0.68% | 1.43% | 0% |
| Logical Memory Immediate Recall | 14.67(3.46) | 9.80(4.25) | 3.87(3.65) |
| Logical Memory Delayed Recall | 13.33(3.69) | 7.39(4.22) | 1.52(3.14) |
| PCA Component 1 | -0.08(0.80) | -0.05(0.87) | 0.64(1.88) |
| PCA Component 2 | -0.05(0.92) | 0.03(1.03) | 0.21(1.35) |
| Motion Parameters (mm) | 0.14 (0.06) | 0.16 (0.07) | 0.17 (0.07) |

### Behavioral Data Processing

Principal Component Analysis (PCA) was used to reduce the dimensionality of this large motor behavioral dataset given that there were multiple measures. We performed the PCA across all the participants with motor variables only, as these variables were dummy coded to indicate ‘normal’ (coded as one) or ‘abnormal’ (coded as two) motor performance by a physician. As a single motor variable coded in that fashion intrinsically reduces variability, using multiple variables via a PCA was intended to increase range and interpretability for our motor measures. Further, this analysis would help us evaluate which measures contributed most to the variability in our sample. Another advantage of the PCA was that it minimized multiple comparisons. This process was not necessary for our cognitive measures which have greater variability by design and had fewer measures. We used jamovi (Version 1.2, retrieved from https://www.jamovi.org) to run the PCA with oblimin rotation and parallel analysis for ADNI dummy coded variables: motor strength, cerebellar finger to nose, cerebellar heel to shin, gait, tremor, plantar reflexes, and deep tendon reflexes on neurological examination.

The Kaiser–Meyer–Olkin (KMO) Measure of Sampling Adequacy and Bartlett’s Test of Sphericity were used to assess the suitability of the respondent data for principal component analysis. The KMO index ranges from 0 to 1, with 0.50 considered suitable for factor analysis (Kaiser, 1970). The Bartlett’s Test of Sphericity should be significant (p < .05) for factor analysis to be suitable (Bartlett, 1950). Direct oblimin rotation was used to determine initial factor structure. Measures were assigned to the component on which they showed the highest loading. We chose two components with the highest loadings.

Composite variables were also created based on PCA component 1 (PCA1) and PCA component 2 (PCA2) factor loadings to verify proof of concept for both PCA component values. These composite variables will be labeled ‘Comp1’ which corresponds with the variables from PCA1 and ‘Comp2’ which corresponds with the variables from PCA2. Comp1 and Comp2 were created by adding the dummy coded values (‘1’ or ‘2’) from each respective PCA component, resulting in a minimum value of 3 or maximum of 6 for each composite variable for a given participant. A greater value would indicate greater motor abnormalities for each participant. The minimum value (3) would indicate no motor abnormalities. The composite variables were created to ensure that our functional analyses were not limited by low variability (i.e., a “floor” value of 3 and a “ceiling” value of 6) and still captured our intent to measure the amount of motor abnormalities. Comparisons of averages by diagnostic group membership are visualized in Figure 1. We did not use the composite variables in the functional imaging analyses. Rather, PCA1 and PCA2 were used in functional analyses to maximize variability for the motor measures in the functional analyses.

**Fig. 1.**
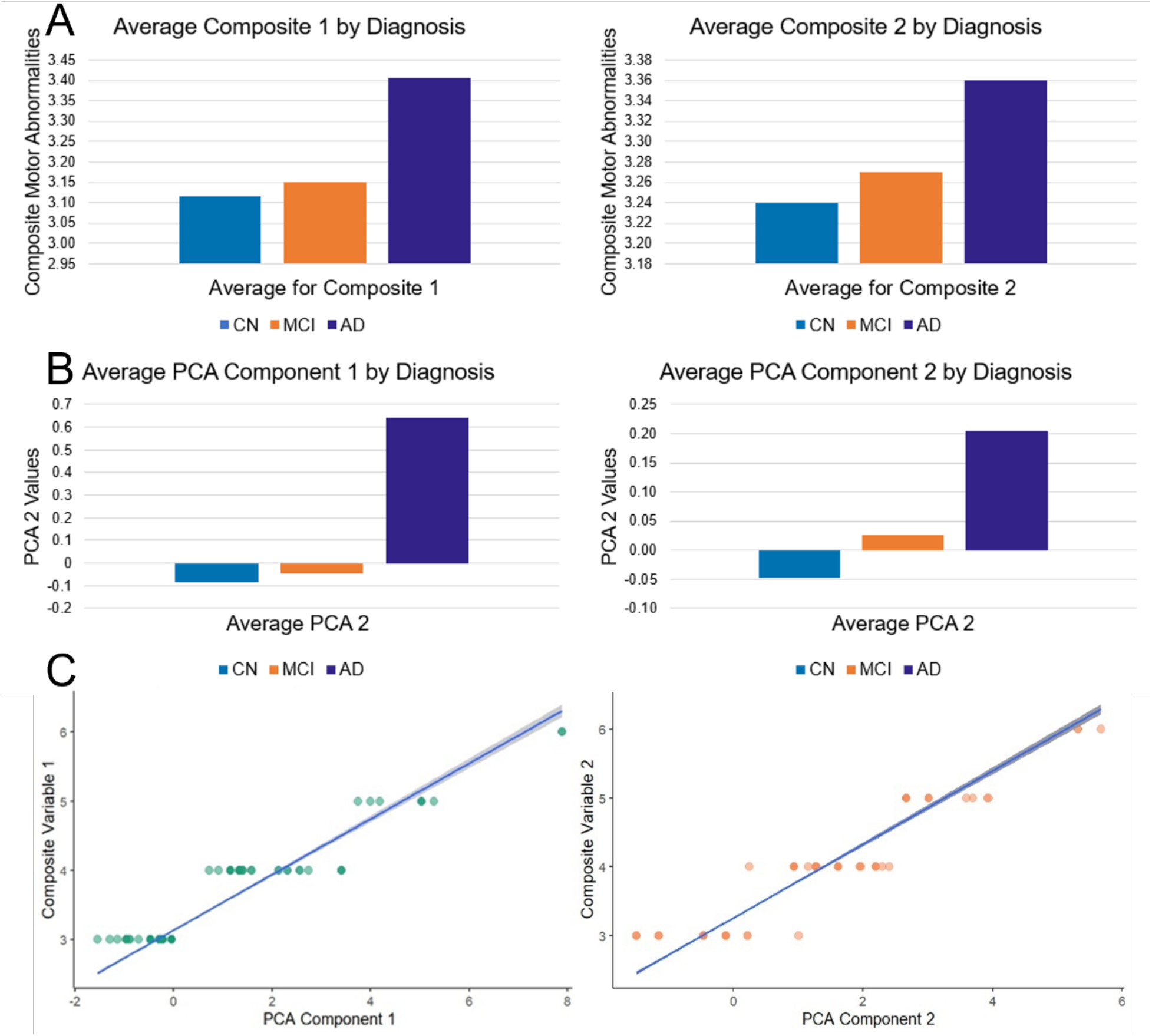
**A**. The top row displays the average composite motor variables (Comp 1, left and Comp 2, right). **Fig 1. B**. The middle row displays the average of the PCA value output (average PCA_*i*_ per the formula above) by diagnostic group (PCA 1, left and PCA 2, right). Comparison of top and bottom rows (right and left sides respectively) exhibit similar patterns by diagnostic group between respective composite scores to their PCA variables. **Fig 1. C**. The scatterplot on the left displays the correlation between PCA Component 1 and Composite variable 1 (*r* (448) = .956) with the gray fill depicting a 95% confidence interval. The scatterplot on the right displays the correlation between PCA Component 2 and Composite variable 2 (*r* (448) = .968) with the gray fill depicting a 95% confidence interval.

For the purposes of this study, subsequent functional imaging analyses were conducted with PCA1 and PCA2 output. The output of the PCA component can be understood as linear combinations of the original variables (e.g., motor strength, tremor, and cerebellar finger to nose) that account for the variance in the data. As our original variables are dummy coded, the numerical PCA output on their own are not easily interpretable. However, these values are still appropriate for this type of analysis. Mathematically, each numerical value output from the PCA per participant would follow this equation, PCA1_*i*_ = ***β***_1_*Motor strength*_*i*_ + ***β***_2_*Tremor*_*i*_ + ***β***_3_*Cerebellar Finger to Nose*_*i*_ + ε_*i*_, where ‘***β***’ (the coefficient) is the result of the linear relationship between each participant’s score and that dummy variable’s relationship to the combined PCA1 variable. The ***β*** coefficients can be calculated in the above formula to produce a PCA1 value for each participant (‘*i*’). The different PCA components elucidate how different aspects of motor functioning (PCA1 versus PCA2) might be loading together in a pattern that could be related to the patterns of neurodegeneration we see in AD. While there is some literature to support certain motor deficits relating to AD and cognitive decline, thus far the type of motor deficits documented have been somewhat inconsistent and nonspecific. This analysis was conceptualized as a first step in parsing out differences that may exist between a wide variety of motor functions.

### Neuroimaging Pre-processing

T1w and BOLD preprocessing were conducted using fMRIPrep version 20.2.3 (20.2.3, 2020). One MPRAGE T1-weighted image was bias and intensity corrected prior to skull stripping, segmentation, and spatial normalization. The BOLD EPI was skull-stripped and corrected for slice time and motion. The EPI was coregistered to a synthetic fieldmap and to the T1w structural image prior to registration to MNI 152 space. Analyses were conducted in standard MNI 152 space. For a detailed description of each preprocessing step, see Supplemental Text 1.

### Statistical Analyses

Preprocessed structural and functional imaging data and head motion, framewise displacement, and confound ROIs (GM, WM, and cerebrospinal fluid (CSF)) were imported into the CONN toolbox (CONN v21a^29^; SPM12^30^; MATLAB^31^), for denoising and analysis. Bandpass filtering was performed prior to time series extraction of each seed (described below). Each seed’s time series was used to create participant-level seed-to-voxel correlation matrices. We calculated seed-to-voxel resting-state functional connectivity using dorsal (MNI; x, y, z: 12, -57, -30) and ventral (MNI; x, y, z: 17, -65, -35) regions of the dentate nucleus defined with 3 mm spherical seeds, selected from previous work.^17,32^

For descriptive statistics in our sample, we used jamovi (Version 1.2) to obtain means and standard deviations for age in years, education in years, mean motion in millimeters (this variable represents the average movement of voxels within an ROI during resting state scans, indicating participant motion in mm during the scan), raw immediate and delayed logical memory scores, and information regarding biological sex. To investigate group differences (AD, MCI, CN) in demographics and behavioral data independent from brain function, we conducted several analyses of variance (ANOVAs) with outcomes: age, education, logical memory immediate and delayed recall (separately), PCA1, and PCA2. Tukey post-hoc analyses were conducted to investigate specific differences between groups. Diagnostic group differences by sex were evaluated via chi-squared independent test of association. Correlations were conducted between each composite variable and the resulting PCA value. Group differences (AD, MCI, CN) in each composite variable were assessed via ANOVAs. Tukey post-hoc analyses were conducted to investigate specific differences between groups.

Our primary analyses modeled dDN and vDN connectivity separately. Due to research showing that females are more likely to develop AD and to experience greater cognitive decline in MCI and AD ^33^, we chose to deviate from the pre-registration and include biological sex in our analyses. Thus, we used a 3 (AD, MCI, CN) x 2 (Male, Female) ANCOVA with education always used as a covariate. Two follow-up analyses were conducted. One explored the main effects of diagnostic grouping using a 3-way ANCOVA (AD, MCI, CN) and the other explored the main effects of sex. Then, group comparisons of diagnostic groupings were completed. Additionally, we were interested in the relationships between dentate connectivity and both motor and cognitive behavior. As such, we modeled group (AD, MCI, CN) x sex (Male, Female) interactions with respect to task performance using the same analysis design. Moreover, we looked at main effects of group and main effects of sex predicting task performance. We followed-up these analyses by conducting group contrasts with respect to task performance. Then, we investigated the relationship between connectivity and behavior in each diagnostic group alone. Finally, for the sake of comparison with past literature on dDN and vDN connectivity, we conducted analyses within each group alone, with CN generally replicating our previous work ^17^; see Supplementary Table 1; Supplementary Figure 1). There were no significant results for AD only, but there were for MCI (Supplementary Table 2). Additionally, a contrast between dDN and vDN networks was run in CN (Supplementary Table 3; Supplementary Figure 2). For all analyses, 1,000 simulations were run using threshold free cluster enhancement with results being *p*<0.05 cluster size p-FDR corrected.

## Results

Table 1 reports basic demographic information and both cognitive (raw scores of correct recalled items in a story, immediate and delayed) and motor performance of the groups. Additionally, ANOVAs by diagnosis covering demographic and motor and cognitive task performance were run. The one-way ANOVA investigating age was significant (F_(2,477)_=9.23, p<.001). Tukey post-hoc analyses revealed significant differences when comparing HC to MCI (t_(2,477)_= -3.25,*p*<.01) and CN to AD (t_(2,477)_= -4.50,*p*<.001), though there were no differences between MCI and AD (t_(2,477)_= -1.49, *p*=0.30). Similarly, there was a significant effect when investigating education (F_(2,477)_=15.14, *p*<.001) and Tukey post-hoc analyses revealed significant differences between both CN and MCI (*p*<.001), and CN and AD (*p*<.001), though the MCI and AD groups did not differ (*p*=0.20). For both the logical memory immediate and delayed recall tests there were significant overall effects (Fs_(2,477)_>207.00, *p*<.001), and all group comparisons were significant (*p*<.001). Chi-squared test of association revealed significant differences in diagnostic group by sex χ^2^(2,477) = 8.73, *p =* .013.

As noted above, because there are multiple motor metrics, and to date, the literature has yet to point to a particular domain as being especially linked to AD, we conducted a PCA to minimize the multiple comparisons and to increase variability. Our assumptions for the PCA were considered suitable: Bartlett’s Test of Sphericity (p<.001) and KMO index = .58. Factor loadings (i.e., Pearson correlation of each item to the component) are displayed in Supplementary Table 4. Component 1 was composed of motor strength, cerebellar finger to nose, and tremor variables which accounted for 23.8% of variance with an eigenvalue of 1.67. Component 2 was composed of gait, plantar reflexes, and deep tendon reflexes which accounted for 16.5% of variance with an eigenvalue of 1.15.

An ANOVA revealed significant group differences for PCA1 (F_(2,477)_=3.26, p=.042). Tukey post-hoc analyses revealed significant group differences on PCA1 between both AD and CN (t(2,477) = -4.61, *p*<.001) and AD and MCI groups (t(2,477) = -4.07,*p*<.001), though the CN and MCI groups did not differ (t(2,477) = -0.357, *p*=.93). With respect to the second PCA component, the ANOVA indicated no significant group effects (F_(2,477)_=.87, *p*=.423). The composite variable based on variables from PCA1, Comp1, was significantly correlated with the values from PCA1 (*r* (448) = .956, *p* <.001). Similarly, the composite variable based on PCA2, Comp2, was significantly correlated with the values from PCA2 (*r* (448) = .968, *p* <.001). As PCA1 is highly correlation with Comp1 and PCA2 is highly correlated with Comp2, we are conceptualizing higher values for both PCA1 and PCA2 as representative of greater motor abnormalities and vice versa with lower values and fewer motor abnormalities. To strengthen our confidence in this conceptualization, we conducted an ANOVA to replicate the findings of our PCA Components with Comp1 and Comp2. This revealed significant group differences for Comp1 (F_(2,477)_=3.55, p=.032). Similarly, Tukey post-hoc analyses replicated significant group differences on Comp1 between both AD and CN (t(2,477) = -4.29, *p*<.001) and AD and MCI groups (t(2,477) = -3.53,*p* = .001), though the CN and MCI groups did not differ (t(2,477) = -0.773, *p*=.72). With respect to Comp2, the ANOVA indicated no significant group effects (F_(2,477)_=.746, *p*=.476). We found nearly identical results in our ANOVA and post-hoc analyses for PCA1 and Comp1 as well as PCA2 and Comp2 (respectively). Overall, our Comp1 and Comp2 analyses indicate that PCA1 and PCA2 values can be interpreted as having a positive linear relationship with the amount of motor abnormalities in these data. As noted in the methods, values from PCA Components 1 and 2 were used for functional imaging analyses for numerical variability.

### Omnibus Models of Connectivity

The 3 (AD, MCI, CN) x 2 (Male, Female) ANCOVA revealed significant group by sex interactions (Supplementary Table 5 and Supplementary Figure 3). From there, we followed up our analyses by running a 3-way ANCOVA exploring the main effects of diagnostic grouping and found significant main effects as well (Supplementary Table 6 and Supplementary Figure 4), though a follow-up test of sex indicated no significant group differences. Finally, we investigated group differences in connectivity (see below).

### Connectivity Differences Between AD and CN

When looking at the dDN seed, we saw higher connectivity in AD relative to CN in the inferior frontal gyrus, thalamus, left angular gyrus, and the middle temporal lobe - areas outside of the traditional motor network of the dentate nucleus^16,17,32^, though there is some extension of these clusters to motor regions (Figures 2). Additionally, there was lower connectivity in AD relative to CN in the middle frontal gyrus, calcarine sulcus, and occipital cortex (Table 2 and Figures 2). For vDN-cortical networks, we found higher FC in the precuneus, caudate, thalamus, and insula in AD relative to the CN group. However, connectivity in AD was lower than CN in the superior frontal gyrus, occipital cortex, right angular gyrus, and cuneus (Supplementary Table 7 and Figures 3). Together, this indicates different patterns of connectivity suggesting that there may be some degree of compensation in AD, though also regions where connectivity was disrupted, relative to the CN group. More broadly, this is indicative of clear differences in DN connectivity patterns in AD.

**Table 2.** Coordinates of the brain regions showing significant differences in FC in the dorsal dentate-cortical network in AD>CN.

| Region (AAL) | X | Y | Z | Cluster Size | T(335) | pFDR |
| --- | --- | --- | --- | --- | --- | --- |
| AD>CN |  |  |  |  |  |  |
| Angular gyrus | -54 | -62 | 40 | 16379 | 4.72 | 0.000034 |
| Middle Temporal gyrus | -62 | -48 | -4 | 2193 | 4.12 | 0.000222 |
| Superior Occipital gyrus | 24 | -88 | 32 | 1798 | 3.84 | 0.000536 |
| Frontal Inferior Orbital gyrus | 42 | 46 | -4 | 174 | 4.68 | 0.000034 |
| Thalamus | -12 | -10 | -2 | 94 | 4.46 | 0.00006 |
| Fusiform gyrus | -34 | -20 | -32 | 53 | 3.47 | 0.001275 |
| Middle Temporal gyrus | -54 | -68 | 6 | 130 | 3.36 | 0.001514 |
| Inferior Temporal gyrus | 62 | -48 | -22 | 18 | 4.74 | 0.000034 |
| Thalamus | 4 | -20 | -4 | 73 | 3.78 | 0.000536 |
| Superior Frontal gyrus | -14 | 18 | 60 | 22 | 3.83 | 0.000536 |
| Opercular Inferior Frontal gyrus | 62 | 14 | 12 | 34 | 3.76 | 0.000536 |
| Vermis I/II | 4 | -40 | -26 | 12 | 3.7 | 0.000625 |
| Cerebellum IV/V | 10 | -52 | -26 | 21 | 3.35 | 0.001514 |
| Inferior Frontal Orbital gyrus | 24 | 34 | -4 | 12 | 4.6 | 0.000038 |
| Cerebellum Crus II | -48 | -58 | -42 | 20 | 2.83 | 0.005507 |
| Middle Temporal gyrus | -54 | -20 | -2 | 13 | 2.72 | 0.0074 |
| Middle Cingulum gyrus | 4 | -14 | 26 | 20 | 2.56 | 0.011137 |
| Vermis I/II | 0 | -34 | -28 | 4 | 3.37 | 0.001514 |
| CN>AD |  |  |  |  |  |  |
| Cerebellum X | 14 | -24 | -48 | 4 | -2.95 | 0.004 |
| Middle Frontal gyrus | -24 | 14 | 32 | 27273 | -4.74 | 0.000034 |
| Calcarine sulcus | 0 | -98 | 2 | 581 | -3.54 | 0.001035 |
| Calcarine sulcus | 18 | -88 | 2 | 118 | -3.23 | 0.001921 |
| Middle Occipital gyrus | -22 | -84 | 6 | 12 | -3.41 | 0.001451 |
| Inferior Occipital gyrus | -28 | -100 | -12 | 15 | -3.03 | 0.003294 |
| Calcarine sulcus | 14 | -106 | 2 | 16 | -3.23 | 0.001921 |
| Superior Frontal gyrus | 20 | 62 | 12 | 26 | -2.83 | 0.005507 |
| Anterior Cingulum cortex | -4 | 42 | 22 | 6 | -3.27 | 0.001834 |
| Middle Frontal gyrus | -32 | 8 | 50 | 20 | -3.27 | 0.001834 |
| Cerebellum VIII | 4 | -74 | -52 | 3 | -3.77 | 0.000536 |
| Middle Occipital gyrus | -34 | -88 | 2 | 4 | -2.54 | 0.01164 |
| Superior Temporal gyrus | 66 | -30 | 8 | 15 | -3.07 | 0.002938 |
| Middle Occipital gyrus | -30 | -102 | -2 | 2 | -3.1 | 0.00279 |

**Fig. 2.**
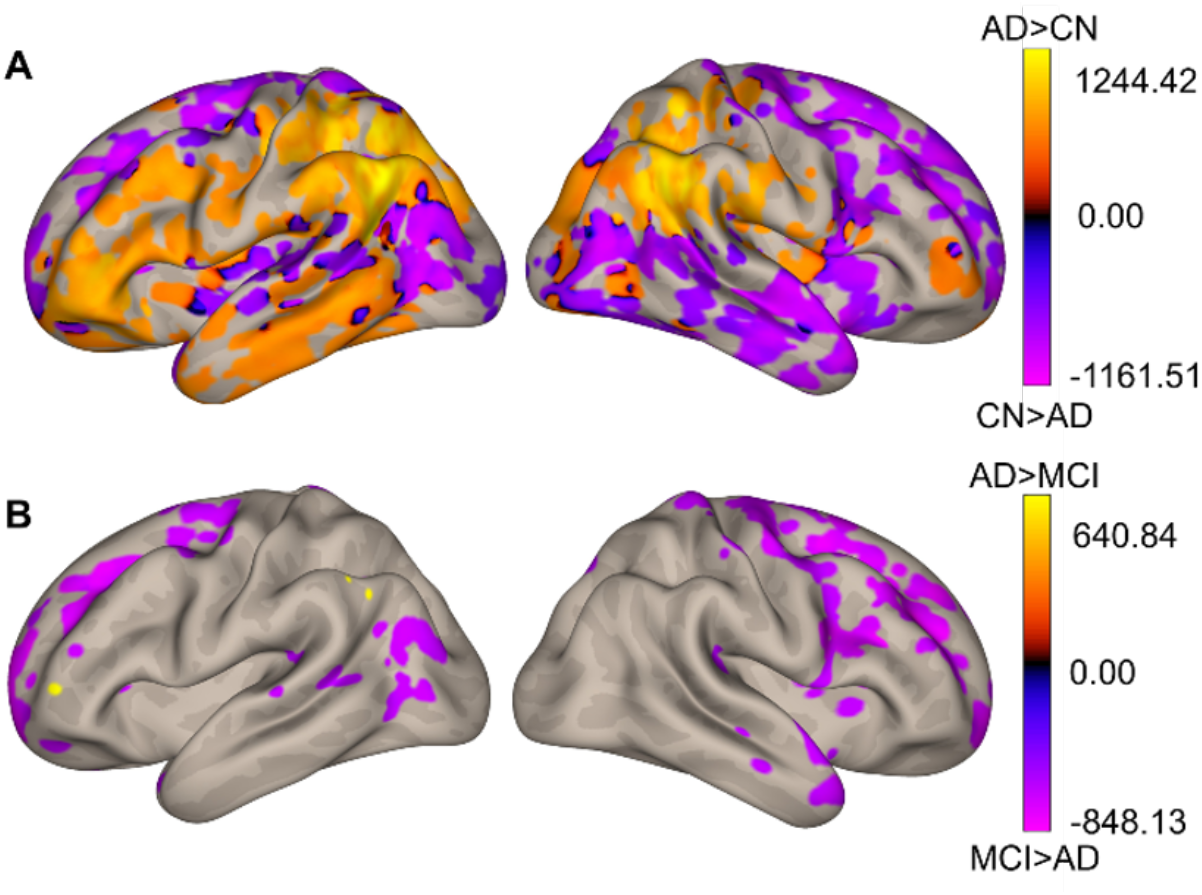
Dorsal Dentate Connectivity. Patterns of differences in FC in the dDN-cortical network. The colorbar represents betas that indicate FC. **A**. orange shows AD>CN and purple represents CN>AD. **B**. orange represents AD>MCI while purple represents MCI>AD.

**Fig. 3.**
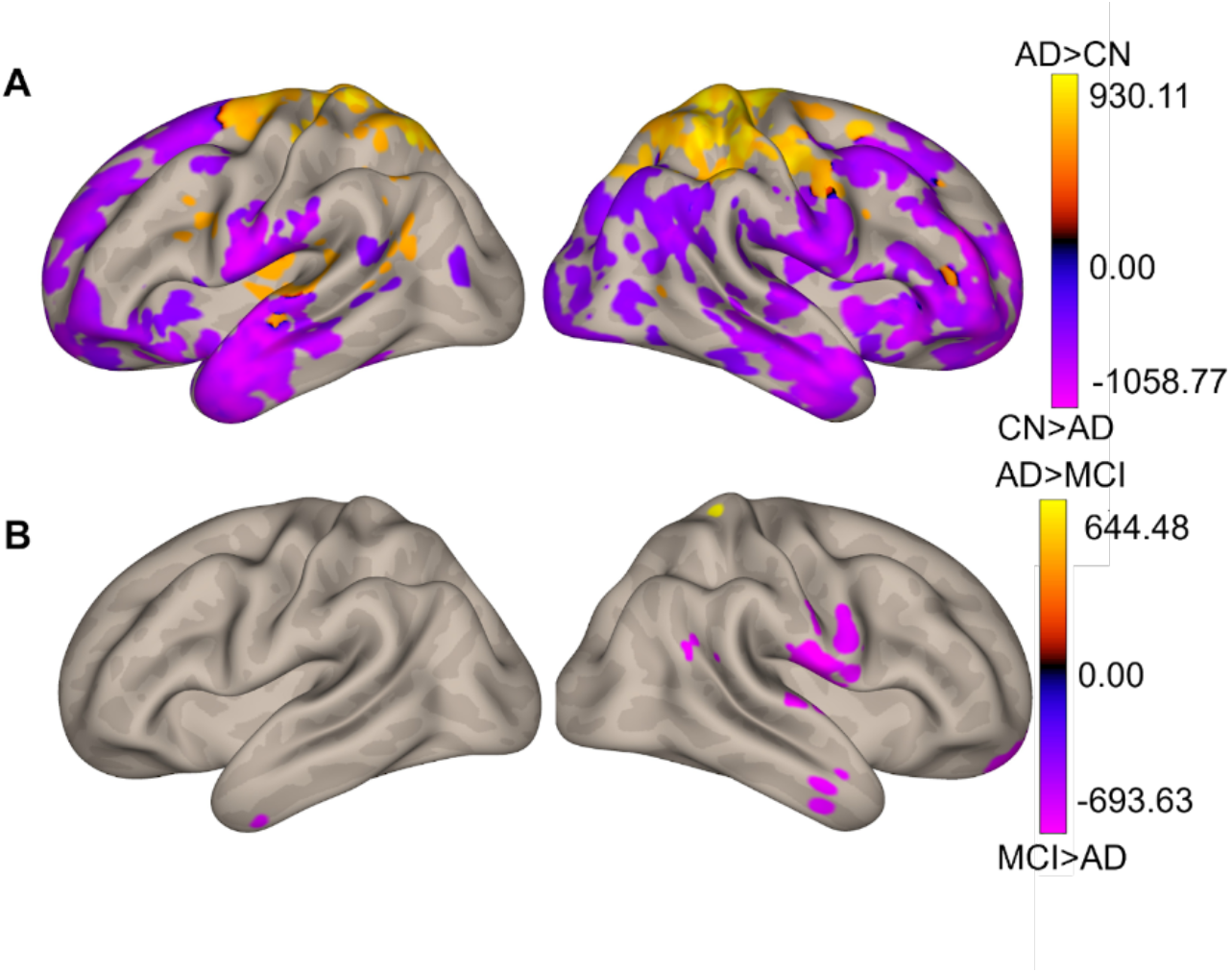
Ventral Dentate Connectivity. Brain regions showing differences in FC in the ventral dentate-cortical network. The color bar represents betas that indicate FC. **A**. Orange indicates AD>CN and purple shows CN>AD. **B**. Orange indicates AD>MCI while purple represents MCI>AD.

### Connectivity Differences Between AD and MCI

When comparing the AD and MCI groups (AD>MCI), we found lower connectivity in AD in the dDN-cortical circuits in areas including the hippocampus and the parahippocampal gyrus, which have been associated with memory encoding and retrieval (Figure 2; Supplementary Table 8). When looking at the vDN, connectivity in AD was lower than that in MCI in the middle frontal gyrus, precuneus, and precentral gyrus (Figure 3; Table 3). Notably, areas where connectivity was greater in the MCI group relative to the AD group show a great deal of spatial overlap with those where connectivity was higher in CN relative to AD (Figures 2 and 4). This may be related to disease progression, suggesting connectivity differs in these regions as individuals progress to more severe disease. Further, across regions where connectivity in AD was lower relative to both CN and MCI, the network in question is largely made up of frontal and temporal regions, while higher connectivity is in distinct areas, regardless of seed. This is indicative of broader connectivity dysfunction, and suggests that this additional network connectivity may be related to compensatory processing. No significant differences were found when comparing the MCI and CN groups, again suggesting more robust differences are associated with more severe disease.

**Table 3.** Coordinates of the brain regions showing differences in FC in the ventral dentate-cortical network in AD>MCI.

| Region | X | Y | Z | Cluster Size | T(182) | pFDR |
| --- | --- | --- | --- | --- | --- | --- |
| AD>MCI |  |  |  |  |  |  |
| Precuneus | 12 | -42 | 64 | 86 | 4.08 | 0.000084 |
| Middle Cingulum gyrus | -12 | -26 | 48 | 74 | 4.17 | 0.000084 |
| Precuneus | -6 | -50 | 60 | 41 | 3.53 | 0.000519 |
| MCI>AD |  |  |  |  |  |  |
| Superior Frontal Orbital cortex | 20 | 56 | -22 | 4 | -4.09 | 0.000084 |
| Superior Temporal cortex | 62 | -6 | 8 | 213 | -4.88 | 0.000012 |
| Superior Frontal Orbital cortex | 14 | 62 | -18 | 66 | -4.91 | 0.000012 |
| Inferior Temporal gyrus | 62 | -6 | -32 | 40 | -4.12 | 0.000084 |
| Postcentral gyrus | 66 | -2 | 20 | 110 | -3.61 | 0.000441 |
| Angular gyrus | 62 | -54 | 26 | 28 | -4.27 | 0.000084 |
| Inferior Temporal gyrus | 46 | -6 | -32 | 22 | -4.22 | 0.000084 |
| Inferior Temporal gyrus | -38 | -4 | -38 | 18 | -4.55 | 0.000036 |

**Fig. 4.**
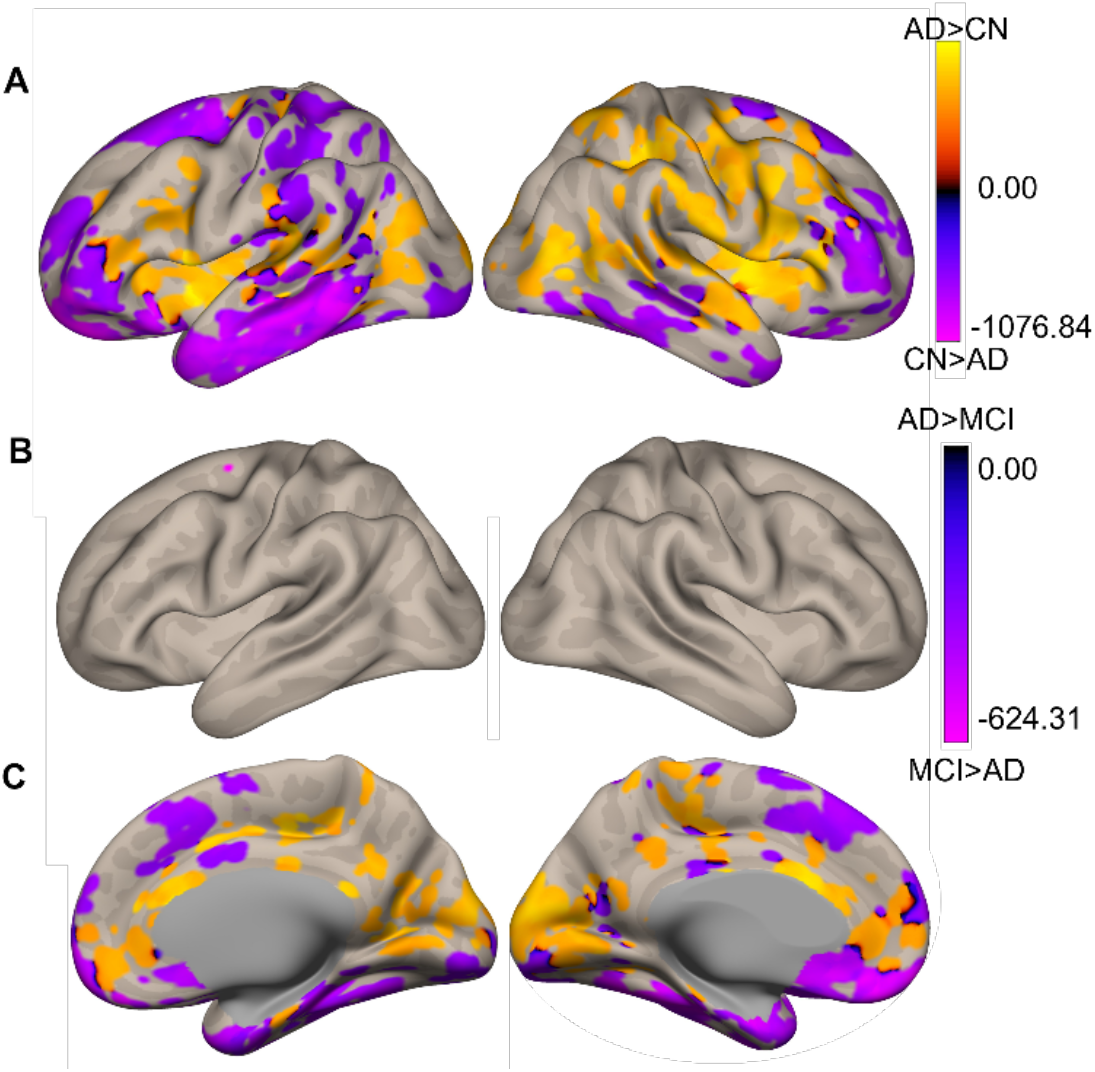
Differences in the relationship between dDN connectivity and immediate recall performance. The color bar represents betas that indicate FC. **A**. orange represents a stronger relationship between dDN connectivity and immediate recall performance in AD relative to CN and purple indicates CN>AD. **B**. black/blue represents AD>MCI and purple shows regions of dDN connectivity where the relationship with immediate recall was stronger in MCI relative to AD. **C**. Medial views of dDN connectivity and immediate recall performance.

### dDN and vDN Connectivity Predict Memory Task Performance

Figure 4 depicts the significant differences in the relationship between FC and performance in an immediate recall task in AD and CN and AD and MCI groups in the dDN-cortical networks, while Figure 5 depicts results for the same interactions in the vDN. (Supplementary Tables 9 and 10; Figures 4 and 5). Here, dDN connectivity in AD is more strongly associated with recall in regions including the superior temporal and middle frontal gyrus as compared to CN. Conversely, connectivity-behavior relationships of the dDN and immediate recall were higher for the CN group in medial aspects of the superior frontal gyrus, middle frontal orbital cortex, and the inferior temporal gyrus compared to AD. Moreover, when looking at the dDN (Supplementary Table 11 and Figure 4), the MCI group had a stronger association between immediate recall and connectivity when compared to AD in areas including the cerebellum lobule VIII and Crus II. On the other hand, in the vdN (Supplementary Table 11 and Figure 5), AD had a stronger relationship compared to MCI in areas including the parahippocampus, the insula, and the temporal pole. No interactions were found for the delayed recall task.

**Fig. 5.**
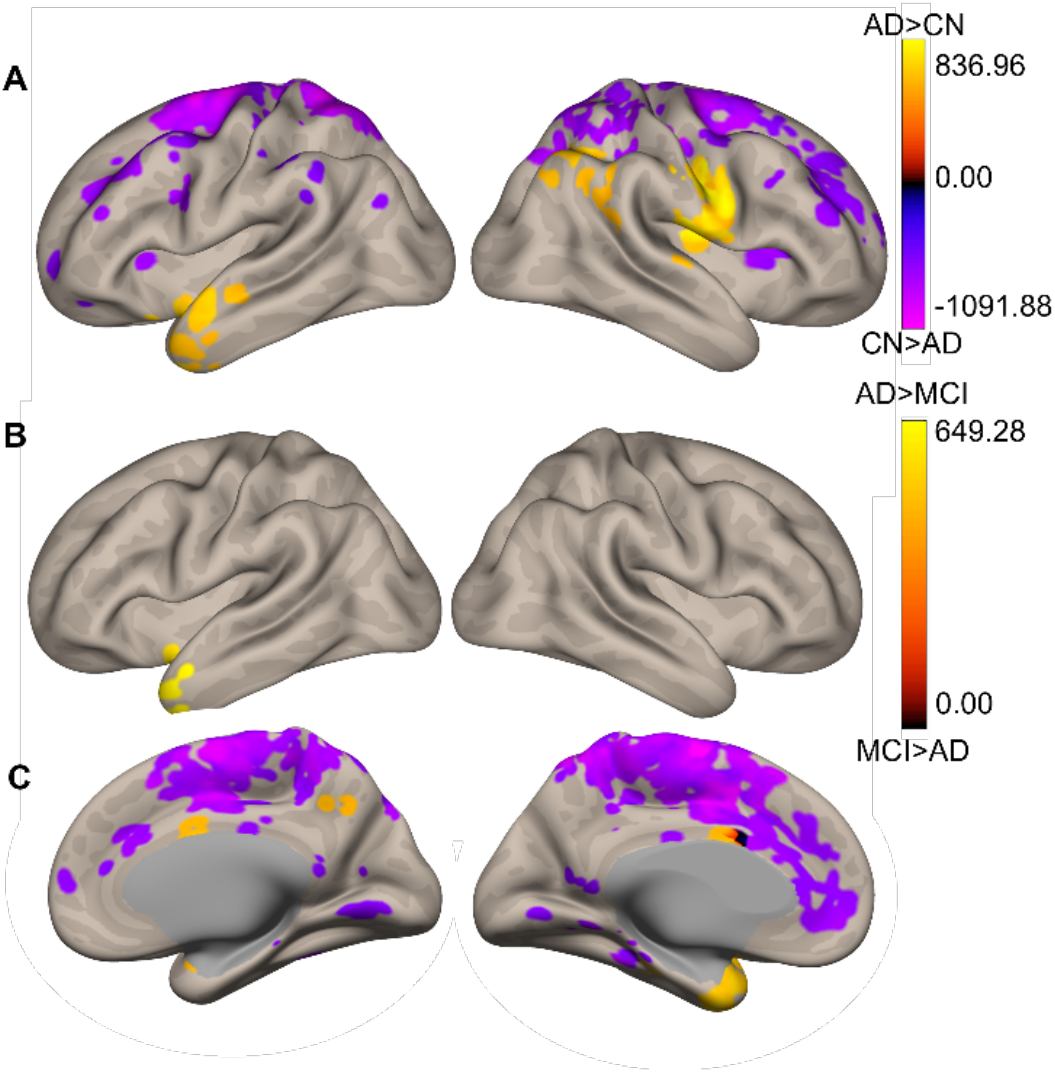
FC connectivity differences in vDN-cortical networks predicting scores for an immediate recall task. The color bar represents betas that indicate FC. In **A**. orange shows AD>CN and purple indicates CN>AD in the relationship associated with immediate recall task performance. In **B**. orange indicates AD>MCI in the relationship between connectivity and task performance while the color black represents MCI>AD. **C**. Medial views of vDN-cortical networks predicting scores for an immediate recall task.

Additionally, there were significant differences in the relationships between FC and motor behavior when investigating PCA 2 (gait, deep tendon and plantar reflexes) between groups in only the dDN-cortical network, suggesting that the dDN is more uniquely associated with motor functions compared to the vDN. These results were found for both AD and MCI relative to CN (Figure 6; Tables 4-5). For example, when looking at the differences in connectivity-behavior relationships between AD and CN, we found that the relationship between connectivity and the motor PCA loading was higher in the precentral gyrus and the temporal pole in AD relative to CN, but when looking at regions where connectivity-behavior associations were strong in CN compared to AD, results were localized to the cingulum cortex and the middle frontal gyrus. No results were found for AD relative to MCI. Additionally, no interactions were found for PCA 1 across any of the three diagnostic groups. Finally, we investigated correlations between connectivity and behavior within each group. We found no significant results except for a relationship in the CN group between PCA 1 and the vDN network, which included the calcarine sulcus and the inferior occipital gyrus (Table 6). Notably however, this was after strict multiple comparisons correction, even after demonstrating significant connectivity-behavior interactions.

**Table 4.**
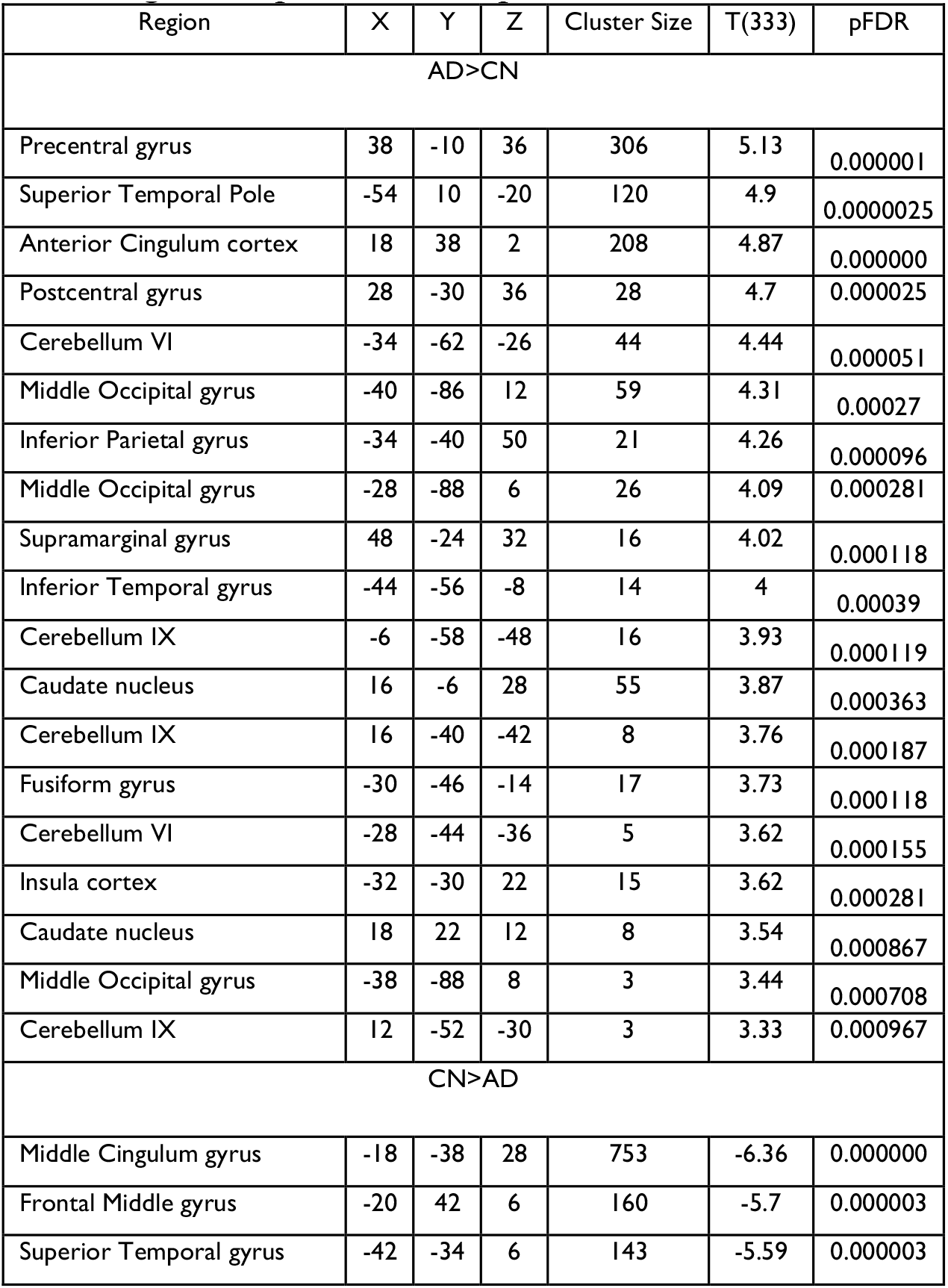
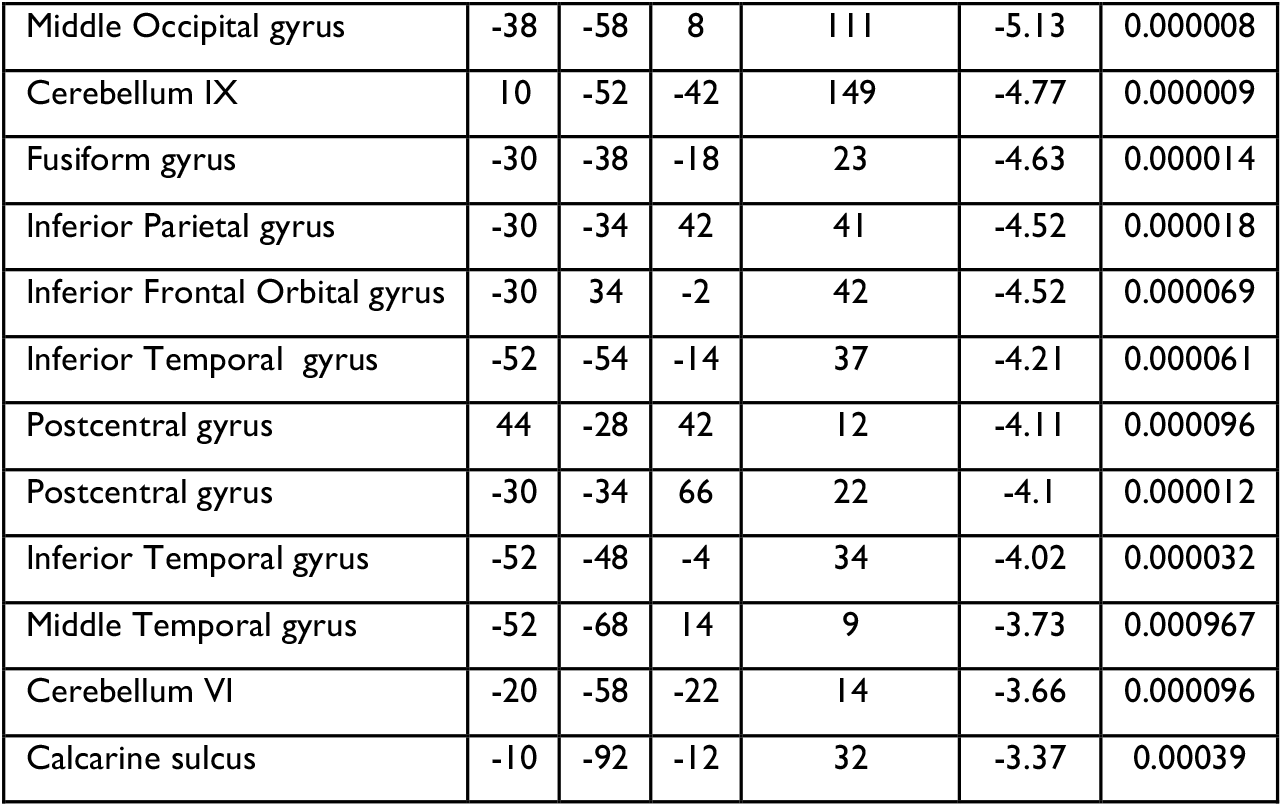
Statistics of FC differences in dDN-cortical networks predicting outcomes in a PCA2 that includes gait, deep tendon, and plantar reflexes in AD>CN.

**Table 5.**
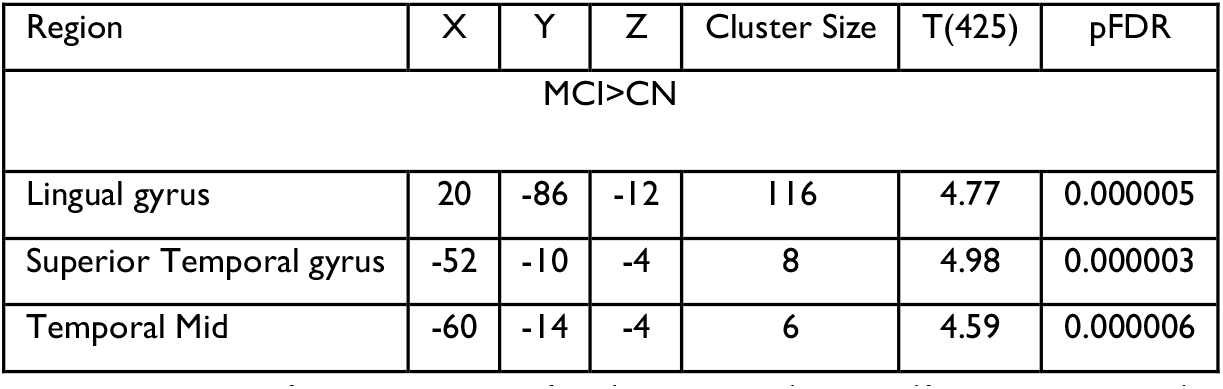
Statistics of FC differences in dDN-cortical networks predicting outcomes in performance of PCA 2 that includes gait, deep tendon, and plantar reflexes in MCI>CN.

**Table 6.**
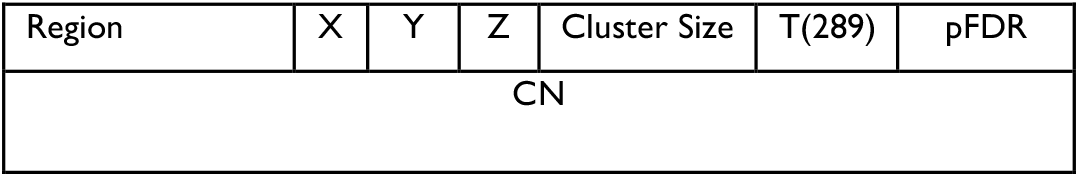
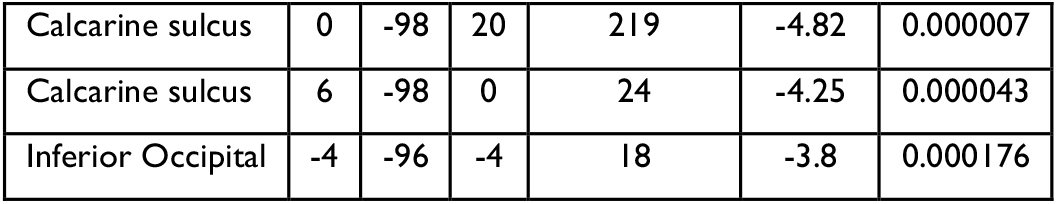
FC in vDN-cortical network predicts motor abnormalities of PCA 1 (tremor, cerebellar finger, and motor strength) in the CN.

**Fig. 6.**
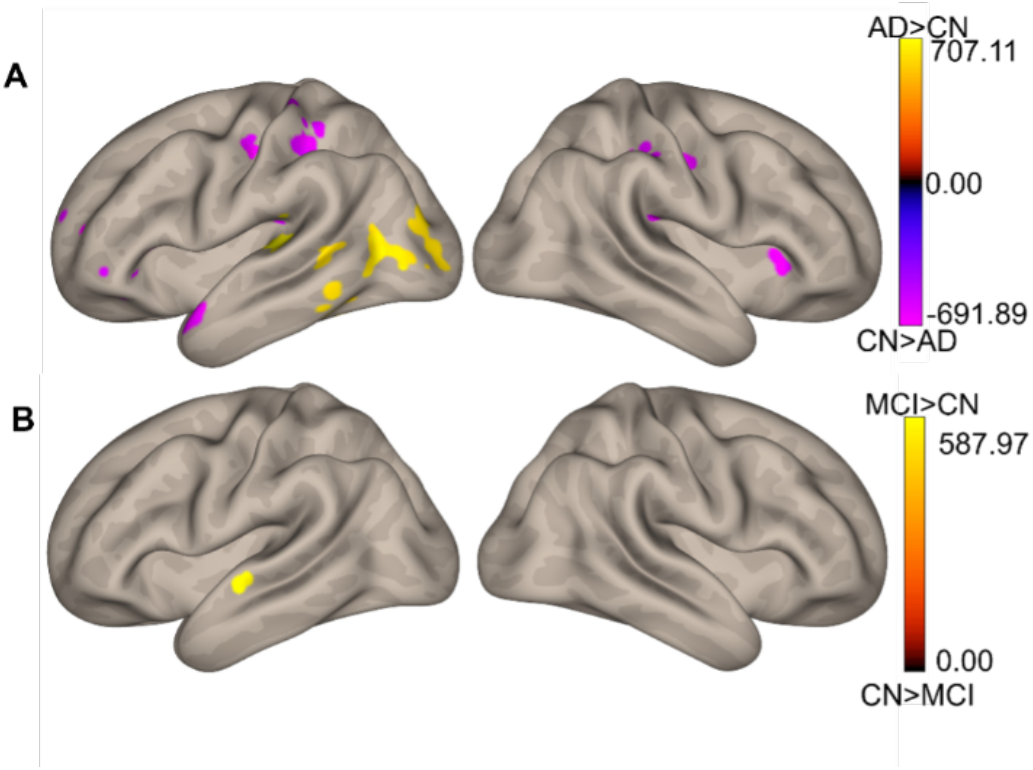
FC differences in dDN-cortical networks predicting outcomes in PCA 2, which includes gait, deep tendon, and plantar reflexes. **A**. orange represents a stronger relationship between dDN connectivity and PCA 2 in AD relative to CN and purple indicates CN>AD. **B**. black represents CN>MCI and purple shows regions of dDN connectivity where the relationship with immediate recall was stronger in MCI relative to CN.

## Discussion

Historically, the cerebellum was thought to be relatively spared in AD. However, a growing literature has shown cerebellar involvement in MCI and AD in the form of GM and WM atrophy.^34,35,36,5,2^ Our findings form part of the growing body of work indicating functional connectivity differences in AD ^37,38,39,40,41^ and relationships with behavioral task performance, including emerging work on cerebellar networks.^26^ Here, we found that the AD group displayed a mix of higher and lower FC in both vDN and dDN networks when compared to CN individuals. When comparing AD to MCI, connectivity was largely lower in the AD group, with some small regions of higher connectivity. Further, cognitive and motor behavior and connectivity are differentially related across diagnostic groups, as evidenced by significant interactions. Motor behavioral classifications were uniquely associated with the dDN, while short term memory was associated with both the dDN and vDN.

Our results here demonstrated that in individuals with AD, there are bidirectional differences in connectivity, when compared to CN and to some degree, MCI groups. Relative to both groups, there are robust patterns of lower connectivity in AD. The cerebellum has been shown to be structurally connected to these regions exhibiting lower connectivity, including the temporal cortex and the hippocampus, which have demonstrated relative atrophy in AD.^42,2^ Since these regions are some of the first ones to exhibit AD pathology, they have been used as biomarkers to track disease progression.^43^ Olivito^26^ demonstrated that besides atrophy and structural connectivity differences, AD affects FC in cerebello-cortical networks. Additionally, it was hypothesized that increased FC may be a compensatory mechanism to mitigate the effects of this disease. While this work was crucial in our understanding of cerebellar connectivity in AD, the present study took an important step forward by exploring MCI as well. While there were differences in AD compared to MCI, with AD showing largely lower FC, we found no significant effects comparing MCI to CN groups. This could suggest that differences in cerebello-cortical FC may be an attempt at compensation in AD, and in MCI there may not yet be a need for cerebellar compensation. Notably however, such compensatory patterns may perhaps be seen in other cortical networks not investigated here. Further, this may relate to disease progression, such that cerebello-cortical network differences are not yet present in those with MCI. As demonstrated through meta-analysis, there are distinct areas of cerebellar atrophy between CN and AD groups^10^, and it may be that in those with MCI, atrophy is more like that in CN, which in turn results in the lack of connectivity differences between MCI and CN described here. Alternatively, it is notable that while an MCI diagnosis (especially amnestic subtype) has a high risk of progressing to AD, it is also possible that someone with MCI will remain stable and even at times revert to a diagnosis of CN^44^ which in turn may also contribute to these findings.

While the areas of lower connectivity may relate to disease progression and pathology, there were also notable regions where connectivity was higher in AD relative to CN, and to some degree MCI. As previously mentioned, we suggest that this may be a potentially compensatory mechanism, or an attempt at compensation in AD, consistent with the broader literature and the assertions of Olivito and colleagues^26^. Research has shown that in healthy aging there is additional cortical activation that may act as a compensatory mechanism to mitigate the effects of aging, described by the compensation-related utilization of neural circuits hypothesis (CRUNCH;^45^). And while CRUNCH focuses on additional activation during task performance, the scaffolding theory of aging and cognition (STAC-R;^46^) defines compensatory scaffolding more broadly. This can include the recruitment of brain networks and/or additional functional brain activation or connectivity. ^47^ Like what we demonstrated here in AD, in Parkinson’s disease (PD), research has shown increased functional connectivity in cerebellar regions. This finding has been interpreted as a reflection of a compensatory mechanism in PD.^48,49,50^ Further work however is necessary to investigate this assertion. We will also note that it is possible that this increased connectivity is not compensatory, but rather an effect of widespread neuropathology and indicative of the disease state in AD.

Regarding behavioral associations with functional connectivity across diagnostic groups, we found relationships that generally highlight the importance of cerebello-cortical networks. In terms of short-term memory as measured by the immediate recall task (for verbal contextual information), the differences in connectivity-behavior relationships were directionally mixed in AD and MCI relative to CN, but not AD relative to MCI. Notably, there was a stronger relationship between FC and immediate recall performance for CN participants in the temporal and frontal gyrus areas as compared to AD. These regions have demonstrated activation during episodic memory tasks on fMRI^51^, indicating that in those with AD, they may be less able to rely on these regions for performance. When looking at the dDN-cortical network, the CN group demonstrated greater FC-behavior relationships than AD in the hippocampus and, similarly, temporal and frontal gyrus and AD connectivity was higher in surrounding areas. This could be interpreted as lower FC in regions responsible for modulating episodic memory tasks due to its associating of worse memory scores in AD, but higher FC in surrounding areas in order to act as compensatory mechanisms for the pathology present. It is notable that we did not see significant FC-behavior relationship differences between the AD and MCI groups when investigating immediate recall. Although performance on this cognitive measure exhibited significant group differences on its own, that data taken alongside FC in the cerebello-cortical networks may not be sensitive or specific enough to parse out differences between earlier versus later stages of AD progression. Alternatively, it may indicate this MCI population includes participants that are less likely convert to AD. Regarding the CN group displaying greater FC than AD in the dDN, it was of particular interest for our study to see these trends in both the hippocampus and parahippocampus. FC in these regions have been associated with episodic memory performance.^52^ Additionally, several cerebellar regions within the dDN-cortical networks demonstrated greater FC in the CN group as compared to AD when predicting scores for the short-term memory task. These findings are fascinating to consider both in the context of cerebellar impact on cognitive networks and AD pathology.

With respect to our PCA2 (containing gait, plantar and deep tendon reflexes) results showed mixed relationships between FC and behavior in AD relative to CN and MCI relative to CN, but not AD to MCI. Koppelmans, Silvester, and Duff’s recent review^53^ found slower gait speed in MCI and AD were associated with lower synchronicity between sensorimotor and frontoparietal networks. While these demonstrate different regions and our results did not reach significance after FDR correction, we did note that CN demonstrated greater FC as compared to AD in dDN to multiple parietal and some frontal networks (i.e., postcentral gyrus, inferior parietal gyrus, and inferior parietal gyrus) in the brain-behavior relationship associated with PCA2. Holtzer and Izzetoglu^54^ revealed declines in prefrontal cortex activation over the course of repeated walking trials in MCI participants using functional near-infrared spectroscopy (fNIRS). If it is common in MCI to have “compensatory” activation and demonstrate significantly reduced activation after several iterations of a cognitive or physical task, this could explain the mixed findings for MCI in the literature and the challenge presented in discriminating between AD and MCI through FC measures whether or not they are interpreted with motor measures that stand to be more sensitive for differentiation at these levels of disease (i.e., MCI to AD). On a more general note, gait speed did predict transition from CN to MCI as well as MCI to unspecified dementia in one longitudinal study.^55^ Although relationships between dentato-cortical networks and PCA2, within group follow-ups, did not demonstrate statistical significance by diagnostic group, PCA1 and PCA2 did demonstrate significant differences between diagnostic groups. It may be that FDR correction is overly stringent, particularly given the significant interactions, and it may be beneficial to further consider these results with more liberal thresholding in new samples. Further understanding cerebello-cortical networks as well as its impact on behavior can elucidate the critical role it plays in mitigating detrimental effects by engaging in compensatory mechanisms.

Historically, the cortex has been the primary focus of AD research. However, investigating networks more broadly, and cerebellar networks specifically, may provide novel insights into neurodegenerative diseases. Much like in AD, research has shown that in schizophrenia (SCZ), pathology is present in the cerebellum and cerebello-thalamo-cortical networks.^56,17,57^ Andreasen^58^ suggested that disruptions to this network were related to the idea of cognitive dysmetria. In addition, Douaud^59^ demonstrated evidence that suggested that AD and SCZ show similar patterns of network deterioration, but at different points of the lifespan. The role of the cerebellum, the similarities in deficits, and the shared pathology underscores the importance of investigating cerebello-cortical networks in neurodegenerative diseases and further highlights the need to incorporate the cerebellum in AD research. Finally, Jacobs and colleagues^2^ summarized the cerebellum’s role in AD, and described how AD pathology can be predicted by the theory of cognitive dysmetria, given that cognitive deficits can be predicted in this context. Our results add to this literature, further emphasizing the cognitive role of the cerebellum and its importance on AD pathology. And while our study was able to find a relationship between connectivity and behavior, it did not explore directionality in terms of task performance and FC differences, which would allow us to understand whether higher or lower FC acts as a compensatory mechanism. Another limitation, due to how motor abnormalities were classified in the ADNI-3 data set, is that we did not explore motor symptom severity in MCI and AD; only if abnormalities were present. Finally, we investigated different stages of cognitive decline cross-sectionally, instead of tracking individual patients longitudinally, which limits our inference related to disease progression.

In the past several decades, canonical views of cerebellar function have shifted to include cognitive and affective roles, in addition to contributions to motor processing. Further, there is an emerging literature in human cognitive neuroscience implicating the cerebellum in AD, though previously it had been thought to be relatively spared. Here, however, we have contributed to an increasing literature that highlights the cerebellum and cerebellar networks’ involvement in AD and MCI. While Olivito and colleagues^26^ provided a foundational understanding of the cerebellar dentate nucleus in AD, their work was limited with the use of one large dentate seed, despite evidence suggesting there are distinct dissociable networks in the dentate nucleus.^16,17^ By investigating the dorsal and ventral dentate separately, we were able to expand our understanding of cerebellar connectivity differences in AD. Further, we showed that deficits in these distinct networks are associated with cognitive and motor behavior in AD. This further highlights that AD symptoms could be due, at least in part, to network deficits consisting of the cerebellum, the thalamus, and the cortex.

## Supporting information

Supplementary Section

## Acknowledgments

*Data used in preparation of this article were obtained from the Alzheimer’s Disease Neuroimaging Initiative (ADNI) database (adni.loni.usc.edu). As such, the investigators within the ADNI contributed to the design and implementation of ADNI and/or provided data but did not participate in analysis or writing of this report. A complete listing of ADNI investigators can be found at: http://adni.loni.usc.edu/wp-content/uploads/how_to_apply/ADNI_Acknowledgement_List.pdf

Data collection and sharing for this project was funded by the Alzheimer’s Disease Neuroimaging Initiative (ADNI) (National Institutes of Health Grant U01 AG024904) and DOD ADNI (Department of Defense award number W81XWH-12-2-0012). ADNI is funded by the National Institute on Aging, the National Institute of Biomedical Imaging and Bioengineering, and through generous contributions from the following: AbbVie, Alzheimer’s Association; Alzheimer’s Drug Discovery Foundation; Araclon Biotech; BioClinica, Inc.; Biogen; Bristol-Myers Squibb Company; CereSpir, Inc.; Cogstate; Eisai Inc.; Elan Pharmaceuticals, Inc.; Eli Lilly and Company; EuroImmun; F. Hoffmann-La Roche Ltd and its affiliated company Genentech, Inc.; Fujirebio; GE Healthcare; IXICO Ltd.; Janssen Alzheimer Immunotherapy Research & Development, LLC.; Johnson & Johnson Pharmaceutical Research & Development LLC.; Lumosity; Lundbeck; Merck & Co., Inc.; Meso Scale Diagnostics, LLC.; NeuroRx Research; Neurotrack Technologies; Novartis Pharmaceuticals Corporation; Pfizer Inc.; Piramal Imaging; Servier; Takeda Pharmaceutical Company; and Transition Therapeutics. The Canadian Institutes of Health Research is providing funds to support ADNI clinical sites in Canada. Private sector contributions are facilitated by the Foundation for the National Institutes of Health (www.fnih.org). The grantee organization is the Northern California Institute for Research and Education, and the study is coordinated by the Alzheimer’s Therapeutic Research Institute at the University of Southern California. ADNI data are disseminated by the Laboratory for Neuro Imaging at the University of Southern California.

This work was further supported by R01AG064010 and R01AG064010-S1 to JAB.

## Notes

### Competing Interest Statement

The authors have declared no competing interest.

