## Supplementary Section for "Functional Connectivity Differences in Distinct Dentato-Cortical Networks in Alzheimer’s Disease and Mild Cognitive Impairment"

**Supplementary Materials**

**Supplementary Text 1.** fMRIPrep boilerplate and citations.

A total of 1 T1-weighted (T1w) images were found within the input Brain Imaging Data Structure (BIDS) dataset. The T1-weighted (T1w) image was corrected for intensity non-uniformity (INU) with N4BiasFieldCorrectioew n (Tustison et al. 2010), distributed with ANTs 2.3.3 (Avants et al. 2008, RRID:SCR_004757), and used as T1w-reference throughout the workflow. The T1w-reference was then skull-stripped with a Nipype implementation of the antsBrainExtraction.sh workflow (from ANTs), using OASIS30ANTs as the target template. Brain tissue segmentation of cerebrospinal fluid (CSF), white matter (WM), and grey matter (GM) were performed on the brain-extracted T1w using fast (FSL 5.0.9, RRID:SCR_002823, Zhang, Brady, and Smith 2001). Volume-based spatial normalization to two standard spaces (MNI152NLin6Asym, MNI152NLin2009cAsym) was performed through nonlinear registration with antsRegistration (ANTs 2.3.3), using brain-extracted versions of both T1w reference and the T1w template. The following templates were selected for spatial normalization: FSL's MNI ICBM 152 non-linear 6th Generation Asymmetric Average Brain Stereotaxic Registration Model [Evans et al. (2012), RRID:SCR_002823; TemplateFlow ID: MNI152NLin6Asym], ICBM 152 Nonlinear Asymmetrical template version 2009c [Fonov et al. (2009), RRID:SCR_008796; TemplateFlow ID: MNI152NLin2009cAsym].

For each of the 1 BOLD runs found per subject (across all tasks and sessions), the following preprocessing was performed. First, a reference volume and its skull-stripped version were generated using a custom methodology of fMRIPrep. A B0-nonuniformity map (or fieldmap) was estimated based on a phase-difference map calculated with a dual-echo GRE (gradient-recall echo) sequence, processed with a custom workflow of SDCFlows inspired by the epidewarp.fsl script and further improvements in HCP Pipelines (Glasser et al. 2013). The fieldmap was then co-registered to the target EPI (echo-planar imaging) reference run and converted to a displacements field map (amenable to registration tools such as ANTs) with FSL's fugue and other SDCflows tools. Based on the estimated susceptibility distortion, a corrected EPI (echo-planar imaging) reference was calculated for a more accurate co-registration with the anatomical reference. The BOLD reference was then co-registered to the T1w reference using flirt (FSL 5.0.9, Jenkinson and Smith 2001) with the boundary-based registration (Greve and Fischl 2009) cost-function. Co-registration was configured with nine degrees of freedom to account for distortions remaining in the BOLD reference. Head-motion parameters with respect to the BOLD reference (transformation matrices, and six corresponding rotation and translation parameters) are estimated before any spatiotemporal filtering using mcflirt (FSL 5.0.9, Jenkinson et al. 2002). BOLD runs were slice-time corrected using 3dTshift from AFNI 20160207 (Cox and Hyde 1997, RRID:SCR_005927). The BOLD time-series (including slice-timing correction when applied) were resampled onto their original, native space by applying a single, composite transform to correct for head-motion and susceptibility distortions. These resampled BOLD time-series will be referred to as preprocessed BOLD in original space, or just preprocessed BOLD. The BOLD time-series were resampled into standard space, generating a preprocessed BOLD run in MNI152NLin6Asym space. First, a reference volume and its skull-stripped version were generated using a custom methodology of fMRIPrep. Several confounding time-series were calculated based on the preprocessed BOLD: framewise displacement (FD), DVARS and three region-wise global signals. FD was computed using two formulations following Power (absolute sum of relative motions, Power et al. (2014)) and Jenkinson (relative root mean square displacement between affines, Jenkinson et al. (2002)). FD and DVARS are calculated for each functional run, both using their implementations in Nipype (following the definitions by Power et al. 2014). The three global signals are extracted within the CSF, the WM, and the whole-brain masks. Additionally, a set of physiological regressors were extracted to allow for component-based noise correction (CompCor, Behzadi et al. 2007). Principal components are estimated after high-pass filtering the preprocessed BOLD time-series (using a discrete cosine filter with 128s cut-off) for the two CompCor variants: temporal (tCompCor) and anatomical (aCompCor). tCompCor components are then calculated from the top 2% variable voxels within the brain mask. For aCompCor, three probabilistic masks (CSF, WM and combined CSF+WM) are generated in anatomical space. The implementation differs from that of Behzadi et al. in that instead of eroding the masks by 2 pixels on BOLD space, the aCompCor masks are subtracted a mask of pixels that likely contain a volume fraction of GM. This mask is obtained by thresholding the corresponding partial volume map at 0.05, and it ensures components are not extracted from voxels containing a minimal fraction of GM. Finally, these masks are resampled into BOLD space and binarized by thresholding at 0.99 (as in the original implementation). Components are also calculated separately within the WM and CSF masks. For each CompCor decomposition, the k components with the largest singular values are retained, such that the retained components' time series are sufficient to explain 50 percent of variance across the nuisance mask (CSF, WM, combined, or temporal). The remaining components are dropped from consideration. The head-motion estimates calculated in the correction step were also placed within the corresponding confounds file. The confound time series derived from head motion estimates and global signals were expanded with the inclusion of temporal derivatives and quadratic terms for each (Satterthwaite et al. 2013). Frames that exceeded a threshold of 0.5 mm FD or 1.5 standardised DVARS were annotated as motion outliers. All resamplings can be performed with a single interpolation step by composing all the pertinent transformations (i.e. head-motion transform matrices, susceptibility distortion correction when available, and co-registrations to anatomical and output spaces). Gridded (volumetric) resamplings were performed using antsApplyTransforms (ANTs), configured with Lanczos interpolation to minimize the smoothing effects of other kernels (Lanczos 1964). Non-gridded (surface) resamplings were performed using mri_vol2surf (FreeSurfer).

**Supplementary Fig. 1** Patterns of differences in FC in healthy controls. The color bar represents betas that indicate FC. **A.** represents FC connectivity patterns associated with the dDN-networks. **B.** indicates FC related to vDN-networks.

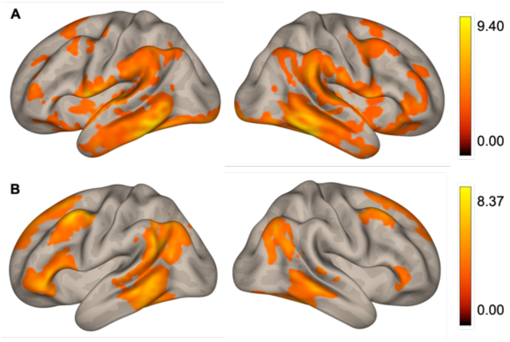

**Supplementary Table 1.** Patterns of differences in FC in CN associated with the dDN- and vDN-cortical networks.

| Region | X | Y | Z | Cluster Size | T(291) | pFDR |
|---|---|---|---|---|---|---|
| **dDN** | | | | | | |
| Cerebellum VIII | 12 | -58 | -30 | 60261 | 26.35 | 0.000000 |
| Supplemental Motor Area | 6 | 8 | 74 | 3351 | 9.68 | 0.000000 |
| Precuneus | 6 | -70 | 44 | 924 | 6.20 | 0.000000 |
| Superior Frontal gyrus | -20 | 22 | 56 | 463 | 7.22 | 0.000000 |
| Middle Frontal gyrus | -24 | 42 | 36 | 143 | 5.36 | 0.000000 |
| Middle Frontal gyrus | -42 | 48 | 8 | 141 | 5.53 | 0.000000 |
| Angular gyrus | 50 | -64 | 46 | 19 | 6.67 | 0.000727 |
| Middle Frontal gyrus | 44 | 24 | 46 | 12 | 3.82 | 0.015123 |
| Postcentral gyrus | 38 | -34 | 66 | 10 | 3.25 | 0.026194 |
| Superior Temporal gyrus | 34 | -34 | 16 | 5 | 4.16 | 0.010818 |
| Supramarginal gyrus | 68 | -16 | 30 | 7 | 5.17 | 0.003846 |
| Supplemental Motor Area | 20 | -20 | 54 | 2 | 3.38 | 0.024159 |
| **vDN** | | | | | | |
| Cerebellum VIII | 18 | -66 | -36 | 34153 | 23.54 | 0.000000 |
| Middle Frontal gyrus | -44 | 6 | 56 | 7920 | 10.40 | 0.000000 |
| Angular gyrus | 48 | -68 | 40 | 846 | 6.15 | 0.000000 |
| Calcarine sulcus | -12 | -94 | -8 | 531 | 4.93 | 0.000001 |
| Inferior Frontal Orbital cortex | 50 | 32 | -2 | 203 | 5.52 | 0.000000 |
| Medial Superior Frontal cortex | -4 | 46 | 40 | 16 | 6.19 | 0.001127 |
| Vermis X | -4 | -28 | -36 | 46 | 3.98 | 0.00805 |
| Opercular Inferior Frontal gyrus | 48 | 22 | 32 | 12 | 4.15 | 0.007603 |
| Superior Medial Frontal gyrus | -12 | 44 | 24 | 49 | 7.1 | 0.000562 |
| Middle Frontal gyrus | 32 | 46 | 2 | 15 | 6.67 | 0.000814 |
| Superior Medial Frontal gyrus | 12 | 32 | 54 | 8 | 4.77 | 0.004217 |
| Postcentral gyrus | 44 | -14 | 32 | 4 | 4.42 | 0.005911 |
| Cerebellum Crus II | -52 | -54 | -42 | 22 | 5.32 | 0.002252 |
| Thalamus | -18 | -28 | 8 | 4 | 5.29 | 0.002252 |
| Superior Frontal gyrus | 20 | 28 | 30 | 2 | 4.1 | 0.007603 |
| Precuneus | 0 | -74 | 50 | 3 | 5.41 | 0.002252 |
| Supramarginal gyrus | 68 | -44 | 30 | 2 | 4.18 | 0.007603 |
| Precuneus | -2 | -78 | 50 | 2 | 5.42 | 0.002252 |
| Supramarginal gyrus | 68 | -44 | 26 | 2 | 3.99 | 0.00805 |
| Superior Medial Frontal gyrus | 12 | 38 | 60 | 1 | 4.49 | 0.005777 |

**Supplementary Table 2.** Patterns of FC differences in MCI only.

| Region | X | Y | Z | Cluster Size | T(137) | pFDR |
|---|---|---|---|---|---|---|
| **dDN** | | | | | | |

| | | | | | | |
|---|---|---|---|---|---|---|
| Cerebellum VI | 18 | -62 | -34 | 192 | 8.35 | 0 |
| **vDN** | | | | | | |
| Cerebellum VIII | 18 | -68 | -40 | 2730.43 | 7.5 | 0 |

**Supplementary Fig. 2** Patterns of differences in FC associated with the contrast between dDN- and vDN-cortical networks in CN only. The color bar represents betas that indicate FC. Orange indicates higher FC in vDN- compared to the dDN-networks while the purple indicates the dDN>vDN contrast.

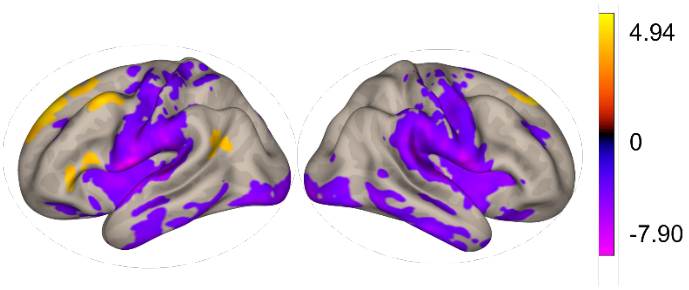


**Supplementary Table 3.** Patterns of differences in FC associated with the contrast between dDN- and vDN-cortical networks.

| Region (AAL) | X | Y | Z | Cluster Size | F(291) | pFDR |
|---|---|---|---|---|---|---|
| **dDN>vDN** | | | | | | |
| Superior Temporal gyrus | 60 | 0 | 6 | 24951 | 15.90 | 0.000000 |
| Fusiform gyrus | 40 | -16 | -28 | 4036 | 11.66 | 0.000000 |
| Cerebellum VIII | 12 | -58 | -30 | 3342 | 16.92 | 0.000000 |
| Cerebellum Crus II | -6 | -88 | -42 | 2917 | 9.68 | 0.000000 |
| Calcarine sulcus | 14 | -68 | 20 | 1885 | 6.48 | 0.000000 |
| Superior Parietal gyrus | -30 | -46 | 58 | 224 | 4.89 | 0.000002 |
| Superior Frontal Orbital cortex | 22 | 34 | -18 | 223 | 6.30 | 0.000000 |
| Middle Frontal gyrus | 32 | 38 | 30 | 186 | 5.34 | 0.000000 |
| Middle Frontal gyrus | -32 | 40 | 32 | 168 | 5.37 | 0.000000 |
| Postcentral gyrus | 20 | -32 | 62 | 91 | 4.72 | 0.000004 |
| Parahippocampal gyrus | 4 | -18 | -20 | 83 | 5.18 | 0.000001 |
| Medial Frontal Orbital cortex | 12 | 44 | 0 | 66 | 4.55 | 0.000008 |
| Precentral gyrus | 30 | -20 | 74 | 55 | 4.38 | 0.000017 |
| Superior Frontal Orbital cortex | 22 | 34 | -18 | 156 | -6.15 | 0.000000 |
| Superior Parietal gyrus | -30 | -46 | 58 | 137 | -4.80 | 0.000000 |
| Middle Frontal gyrus | 32 | 38 | 30 | 131 | -5.38 | 0.000000 |
| Vermis IV/V | 00 | -52 | 06 | 130 | 5.76 | 0.000000 |
| Cerebellum Crus I | -44 | -40 | -38 | 130 | 6.25 | 0.000000 |
| Inferior Frontal Orbital cortex | -42 | 36 | -2 | 127 | 5.35 | 0.000000 |
| Middle Frontal gyrus | -32 | 40 | 32 | 124 | -5.23 | 0.000000 |

| | | | | | | |
|---|---|---|---|---|---|---|
| Inferior Parietal gyrus | 54 | -36 | 54 | 99 | -5.31 | 0.000000 |
| Middle Frontal Orbital cortex | -28 | 38 | -14 | 96 | -5.46 | 0.000000 |
| Posterior Cingulum cortex | -4 | -28 | 28 | 88 | -5 | 0.000001 |
| Parahippocampal gyrus | 04 | -18 | -20 | 61 | -5.11 | 0.000001 |
| Postcentral gyrus | 20 | -32 | 62 | 58 | -4.83 | 0.000003 |
| Medial Frontal_Orbital_cortex | 12 | 44 | 0 | 56 | -4.57 | 0.000008 |
| Postcentral gyrus | 40 | -28 | 58 | 52 | -4.25 | 0.000029 |
| Precentral gyrus | 30 | -20 | 74 | 43 | -4.48 | 0.000011 |
| Inferior Frontal_Operculum | -60 | 18 | 16 | 42 | 4.41 | 0.000015 |
| **vDN>dDN** | | | | | | |
| Cerebellum VIII | 18 | -66 | -36 | 5767 | -19.03 | 0.000000 |
| Superior Frontal gyrus | -16 | 32 | 60 | 1195 | -6.98 | 0.000000 |
| Hippocampus | 34 | -40 | 2 | 681 | -7.75 | 0.000000 |
| Insula cortex | -26 | -36 | 24 | 562 | -6.90 | 0.000000 |
| Precentral gyrus | -42 | 4 | 62 | 253 | -5.84 | 0.000000 |
| Angular gyrus | -48 | -58 | 26 | 234 | -5.26 | 0.000000 |
| Inferior Frontal Orbital cortex | 42 | 36 | -2 | 214 | -5.35 | 0.000000 |
| Vermis IV/V | 0 | -52 | 6 | 157 | -5.77 | 0.000000 |

**Supplementary Table 4.** Principal Component Analysis Factor Loadings

| | Component 1 | Component 2 |
|---|---|---|
| Motor Strength | 0.578 | |
| Tremor | 0.670 | |
| Cerebellar Finger-Nose | 0.765 | |
| Gait | | 0.728 |
| Plantar Reflexes | | 0.559 |
| Deep Tendon | | 0.657 |

**Supplementary Table 5.** 3x2 ANCOVA: Diagnostic Grouping x Sex of both dDN- and vDN-cortical networks.

| Region | X | Y | Z | Cluster Size | T(3470) | pFDR |
|---|---|---|---|---|---|---|
| **dDN** | | | | | | |
| Lingual gyrus | 10 | -58 | 8 | 49457 | 2.92 | 0.036856 |
| Precentral | 34 | -26 | 60 | 284 | 4.21 | 0.010818 |
| Superior Frontal Orbital Cortex | -14 | 28 | -30 | 70 | 6.54 | 0.000727 |
| Cerebellum Crus II | -34 | -72 | -48 | 92 | 6.6 | 0.000727 |
| Cerebellum Crus II | 44 | -72 | -48 | 31 | 7.62 | 0.000659 |
| Middle Frontal gyrus | 28 | 38 | 42 | 49 | 1.8 | 0.146816 |
| Angular gyrus | 50 | -64 | 46 | 19 | 6.67 | 0.000727 |
| Middle Frontal gyrus | 44 | 24 | 46 | 12 | 3.82 | 0.015123 |
| Postcentral gyrus | 38 | -34 | 66 | 10 | 3.25 | 0.026194 |
| Superior Temporal gyrus | 34 | -34 | 16 | 5 | 4.16 | 0.010818 |
| Supramarginal gyrus | 68 | -16 | 30 | 7 | 5.17 | 0.003846 |
| Supplementary Motor Area | 20 | -20 | 54 | 2 | 3.38 | 0.024159 |
| **vDN** | | | | | | |
| Paracentral Lobule | -8 | -36 | 64 | 13728 | 8.17 | 0.000246 |
| Cerebellum IV/V | 18 | -36 | -22 | 4969 | 9.64 | 0.000069 |
| Thalamus | -24 | -30 | 12 | 47 | 7.91 | 0.000246 |
| Superior Frontal Medial gyrus | -4 | 32 | 60 | 73 | 5.96 | 0.001344 |
| Middle Frontal gyrus | 24 | 52 | 2 | 48 | 6.29 | 0.001127 |
| Superior Frontal Medial gyrus | -4 | 46 | 40 | 16 | 6.19 | 0.001127 |
| Vermis X | -4 | -28 | -36 | 46 | 3.98 | 0.00805 |
| Inferior Frontal Operculum | 48 | 22 | 32 | 12 | 4.15 | 0.007603 |
| Superior Frontal Medial gyrus | -12 | 44 | 24 | 49 | 7.1 | 0.000562 |
| Middle Frontal gyrus | 32 | 46 | 2 | 15 | 6.67 | 0.000814 |
| Superior Frontal Medial gyrus | 12 | 32 | 54 | 8 | 4.77 | 0.004217 |
| Postcentral gyrus | 44 | -14 | 32 | 4 | 4.42 | 0.005911 |
| Cerebellum Crus II | -52 | -54 | -42 | 22 | 5.32 | 0.002252 |
| Thalamus | -18 | -28 | 8 | 4 | 5.29 | 0.002252 |
| Superior Frontal gyrus | 20 | 28 | 30 | 2 | 4.1 | 0.007603 |
| Precuneus | 0 | -74 | 50 | 3 | 5.41 | 0.002252 |
| Supramarginal gyrus | 68 | -44 | 30 | 2 | 4.18 | 0.007603 |
| Precuneus | -2 | -78 | 50 | 2 | 5.42 | 0.002252 |
| Supramarginal gyrus | 68 | -44 | 26 | 2 | 3.99 | 0.00805 |
| Superior Frontal Medial gyrus | 12 | 38 | 60 | 1 | 4.49 | 0.005777 |

**Supplementary Fig. 3.** 3x2 ANCOVA: Diagnostic Grouping x Sex of both dDN and vDN networks. The color bar represents betas that indicate FC. **A.** represents the interactive effect of Dx x Sex in dDN-cortical networks. **B.** indicates the interaction of Dx x Sex in vDN-cortical networks.

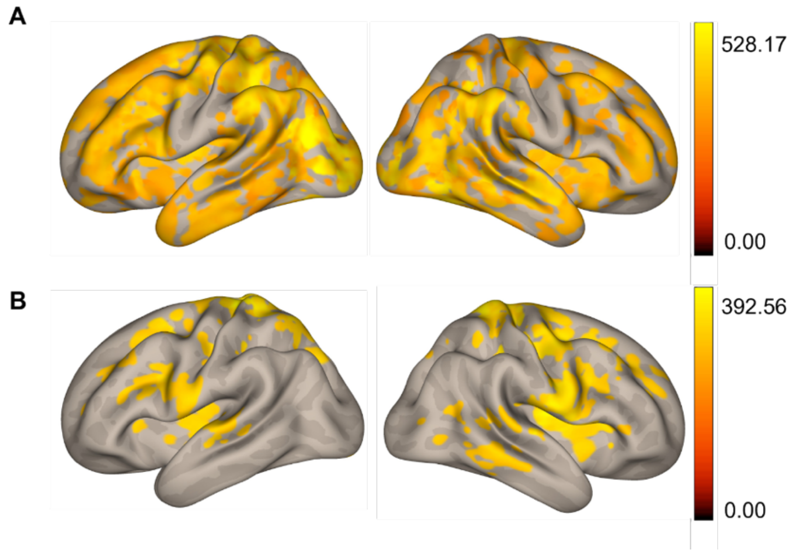


**Supplementary Table 6.** Main Effects of Diagnostic Grouping of both dDN- and vDN-cortical networks.

| Region (AAL) | X | Y | Z | Cluster Size | F([2 474]) | pFDR |
|---|---|---|---|---|---|---|
| **dDN** | | | | | | |
| Cerebellum IX | 6 | -40 | -38 | 445 | 16.77 | 0.000002 |
| Precentral gyrus | 24 | -14 | 42 | 403 | 18.33 | 0.000001 |
| Lingual gyrus | 20 | -64 | -4 | 825 | 13.82 | 0.00002 |
| Middle Occipital gyrus | -36 | -68 | 26 | 1777 | 8.31 | 0.000682 |
| Caudate nucleus | -20 | -18 | 30 | 612 | 7.11 | 0.001613 |
| Caudate nucleus | -14 | 18 | 8 | 289 | 12.6 | 0.000048 |
| Superior Occipital gyrus | 18 | -78 | 36 | 808 | 9.86 | 0.000218 |
| Fusiform gyrus | 38 | -14 | -26 | 130 | 11.17 | 0.000083 |
| Lingual gyrus | -6 | -82 | -4 | 280 | 6.4 | 0.002661 |
| Lingual gyrus | -18 | -54 | -2 | 179 | 11.13 | 0.000083 |
| Inferior Temporal gyrus | -40 | -14 | -24 | 101 | 11.83 | 0.000071 |
| Inferior Frontal Orbital cortex | 32 | 24 | -22 | 66 | 11.75 | 0.000071 |
| Middle Cingulum | 6 | -34 | 40 | 32 | 11.11 | 0.000083 |
| Superior Frontal gyrus | 14 | 42 | 38 | 103 | 8.53 | 0.000588 |
| Lingual gyrus | -18 | -78 | -12 | 59 | 7.87 | 0.000994 |
| Caudate nucleus | -16 | 16 | 26 | 34 | 4.17 | 0.016066 |
| Middle Frontal gyrus | 24 | 42 | 8 | 14 | 11.05 | 0.000083 |
| Vermis I/II | 0 | -30 | -26 | 23 | 9.65 | 0.000245 |
| Temporal Sup | -58 | -16 | 6 | 61 | 6.66 | 0.002138 |
| Cerebellum X | -6 | -30 | -42 | 14 | 10.59 | 0.000118 |
| Postcentral gyrus | -58 | -8 | 38 | 53 | 5.85 | 0.003734 |
| Insula cortex | -28 | 22 | -4 | 17 | 9.07 | 0.000372 |

| | | | | | | |
|---|---|---|---|---|---|---|
| Anterior Cingulum | 14 | 34 | 22 | 23 | 7.27 | 0.001441 |
| Middle Cingulum | -20 | -34 | 46 | 22 | 6.03 | 0.00321 |
| Caudate nucleus | 14 | 4 | 12 | 30 | 7.48 | 0.001297 |
| Caudate nucleus | 18 | 28 | 8 | 26 | 7.64 | 0.001166 |
| Lingual gyrus | -20 | -54 | -12 | 4 | 6.3 | 0.002815 |
| Middle Temporal gyrus | 50 | -10 | -18 | 51 | 7.36 | 0.001384 |
| Vermis IV /V | 0 | -52 | -4 | 7 | 7.05 | 0.001643 |
| Middle Frontal gyrus | -22 | 22 | 28 | 14 | 6.11 | 0.003085 |
| Superior Temporal gyrus | -52 | -14 | -4 | 8 | 5.4 | 0.005324 |
| Rolandic Operculum | -52 | -2 | 6 | 7 | 6.11 | 0.003085 |
| Caudate nucleus | -20 | 26 | 22 | 12 | 5.44 | 0.00525 |
| Cerebellum III | -10 | -28 | -30 | 3 | 6.72 | 0.002093 |
| Middle Temporal Pole | 36 | 16 | -32 | 15 | 9.14 | 0.000372 |
| Middle Cingulum | -10 | -30 | 46 | 13 | 5.44 | 0.00525 |
| Superior Temporal gyrus | -44 | -10 | -4 | 4 | 4.42 | 0.012795 |
| Cerebellum VI | 34 | -34 | -32 | 20 | 6.77 | 0.00207 |
| Cerebellum IV/V | 24 | -44 | -18 | 12 | 5.13 | 0.006771 |
| Middle Occipital gyrus | -38 | -68 | 38 | 4 | 4.77 | 0.009375 |
| Cerebellum III | -8 | -32 | -26 | 2 | 6.11 | 0.003085 |
| **vDN** | | | | | | |
| Insula cortex | -30 | 14 | -20 | 581 | 16.69 | 0.000002 |
| Angular gyrus | 48 | -66 | 38 | 78 | 14.1 | 0.000007 |
| Fusiform gyrus | -28 | -8 | -36 | 222 | 14.52 | 0.000007 |
| Middle Cingulum | 14 | -30 | 26 | 51 | 13.55 | 0.000009 |
| Pallidum | 14 | 8 | 2 | 43 | 12.89 | 0.000013 |
| Lingual gyrus | 24 | -52 | -4 | 105 | 12.83 | 0.000013 |
| Caudate nucleus | -18 | 22 | 0 | 71 | 9.39 | 0.000185 |
| Pallidum | -14 | 6 | 2 | 36 | 14.18 | 0.000007 |
| Superior Frontal gyrus | 32 | 58 | 12 | 59 | 9.53 | 0.000175 |
| Middle Cingulum | 20 | -30 | 40 | 17 | 11.79 | 0.00003 |
| Superior Frontal gyrus | 28 | 46 | 22 | 98 | 8.31 | 0.000401 |
| Insula cortex | 44 | 14 | -10 | 37 | 8.37 | 0.000401 |
| Middle Temporal Pole | -42 | 12 | -44 | 56 | 7.96 | 0.000531 |
| Fusiform gyrus | 32 | -10 | -38 | 69 | 11.52 | 0.000035 |
| Parahippocampal gyrus | 10 | -6 | -36 | 16 | 10.02 | 0.000119 |
| Middle Frontal Orbital cortex | -20 | 42 | -18 | 16 | 10.14 | 0.000117 |
| Frontal Sup | 28 | 48 | 12 | 24 | 8.73 | 0.000323 |
| Frontal Med Orb | -2 | 40 | -10 | 24 | 8.36 | 0.000401 |
| Pallidum | 14 | 8 | -2 | 2 | 7.35 | 0.000818 |
| Temporal Mid | -40 | -2 | -26 | 1 | 4.92 | 0.008 |
| Putamen | 18 | 18 | -2 | 8 | 7.21 | 0.0009 |
| Caudate | -10 | 12 | 2 | 2 | 7.44 | 0.000792 |

| | | | | | | |
|---|---|---|---|---|---|---|
| Putamen | -24 | -2 | 8 | 4 | 7.75 | 0.000615 |
| Fusiform | -32 | 2 | -32 | 1 | 4.7 | 0.009483 |

**Supplementary Table 7.** Coordinates of the brain regions showing significant differences in FC in the ventral dentate-cortical network in AD>CN.

| Region | X | Y | Z | Cluster Size | T(335) | pFDR |
|---|---|---|---|---|---|---|
| AD>CN | | | | | | |
| Precuneus | -6 | -52 | 64 | 5162 | 4.16 | 0.000309 |
| Caudate nucleus | 18 | 4 | 30 | 87 | 4.34 | 0.000256 |
| Insula cortex | -34 | -10 | 26 | 90 | 4.04 | 0.000309 |
| Superior Temporal gyrus | -54 | -24 | 10 | 277 | 4.47 | 0.000227 |
| Middle Occipital gyrus | -30 | -64 | 32 | 60 | 3.74 | 0.000541 |
| Anterior Cingulum cortex | -6 | 4 | 30 | 99 | 4.14 | 0.000309 |
| Inferior Temporal gyrus | 42 | -38 | -8 | 42 | 4.17 | 0.000309 |
| Lingual gyrus | 14 | -28 | -4 | 102 | 3.95 | 0.000378 |
| Superior Frontal gyrus | 28 | -10 | 66 | 38 | 3.57 | 0.000728 |
| Putamen | -30 | -20 | -2 | 54 | 3.65 | 0.00064 |
| Vermis IV/V | 4 | -54 | 6 | 22 | 4.14 | 0.000309 |
| Middle Cingulum cortex | -20 | -38 | 40 | 24 | 4.07 | 0.000309 |
| Pars Triangularis of the Inferior Frontal gyrus | 42 | 32 | 6 | 27 | 4.08 | 0.000309 |
| Thalamus | -24 | -28 | 16 | 37 | 3.72 | 0.000541 |
| Putamen | 30 | -8 | 6 | 36 | 4.08 | 0.000309 |
| Hippocampus | 24 | -40 | 12 | 20 | 4.05 | 0.000309 |
| Thalamus | -24 | -24 | 2 | 11 | 3.66 | 0.00064 |
| Rolandic Operculum | -40 | -20 | 16 | 44 | 3.39 | 0.001108 |
| Middle Temporal gyrus | -38 | -48 | 6 | 12 | 3.72 | 0.000541 |
| Anterior Cingulum cortex | -18 | 32 | 20 | 17 | 3.58 | 0.000728 |
| Anterior Cingulum cortex | 18 | 34 | 26 | 42 | 3.56 | 0.000728 |
| Putamen | 28 | 8 | 6 | 16 | 3.97 | 0.000355 |
| Caudate nucleus | -20 | 0 | 30 | 21 | 3.35 | 0.00125 |
| Thalamus | 10 | -16 | 20 | 10 | 3.83 | 0.000502 |
| Superior Temporal gyrus | -48 | -20 | 12 | 17 | 3.21 | 0.001682 |
| Lingual gyrus | 26 | -52 | -2 | 21 | 3.64 | 0.00064 |
| Cerebellum Crus I | 34 | -72 | -32 | 14 | 3.76 | 0.000541 |
| Inferior Frontal Operculum | 34 | 4 | 26 | 4 | 3.57 | 0.000728 |
| Middle Temporal gyrus | -44 | -62 | 22 | 17 | 3.28 | 0.001397 |
| Inferior Frontal Operculum | -44 | 8 | 30 | 16 | 3.62 | 0.000657 |
| Middle Occipital gyrus | -38 | -62 | 22 | 8 | 3.34 | 0.00125 |
| Superior Temporal gyrus | -62 | -28 | 12 | 8 | 3.04 | 0.00275 |
| Caudate nucleus | 20 | 24 | 22 | 22 | 3.69 | 0.000579 |
| Putamen | 28 | -22 | 10 | 13 | 3.64 | 0.00064 |
| Middle Temporal gyrus | 48 | -44 | -4 | 24 | 3.53 | 0.000815 |

| | | | | | | |
|---|---|---|---|---|---|---|
| Insula cortex | -24 | 24 | 12 | 17 | 3.33 | 0.001262 |
| Middle Temporal gyrus | -44 | -48 | 6 | 4 | 3.51 | 0.000836 |
| Middle Frontal Orbital cortex | 34 | 42 | -8 | 4 | 3.73 | 0.000541 |
| Middle Temporal gyrus | -42 | -52 | 12 | 4 | 3.46 | 0.000952 |
| Insula cortex | -38 | -16 | 2 | 12 | 3.45 | 0.000954 |
| Pars Triangularis of the Inferior Frontal gyrus | -44 | 14 | 26 | 8 | 3.73 | 0.000541 |
| Postcentral gyrus | -42 | -34 | 64 | 4 | 2.86 | 0.004703 |
| Middle Cingulum cortex | 10 | 8 | 36 | 4 | 3.32 | 0.001301 |
| Superior Frontal gyrus | 18 | -14 | 60 | 10 | 2.92 | 0.004014 |
| Insula cortex | 28 | 18 | 16 | 8 | 3.3 | 0.001353 |
| Putamen | 30 | 4 | 16 | 11 | 3.64 | 0.00064 |
| Inferior Frontal Orbital cortex | 38 | 36 | -2 | 1 | 3.43 | 0.000995 |
| Angular gyrus | -42 | -64 | 26 | 1 | 3.17 | 0.001903 |
| Caudate nucleus | -14 | 22 | 22 | 4 | 3.23 | 0.0016 |
| Inferior Frontal Orbital cortex | 36 | 34 | 0 | 1 | 3.29 | 0.001364 |
| Thalamus | 22 | -26 | 12 | 2 | 3.28 | 0.001397 |
| Superior Parietal gyrus | 18 | -48 | 56 | 2 | 2.42 | 0.016207 |
| CN>AD | | | | | | |
| Superior Frontal Orbital cortex | 14 | 62 | -18 | 19937 | -5.29 | 0.00002 |
| Inferior Occipital cortex | 30 | -86 | -12 | 2097 | -5.15 | 0.00002 |
| Angular gyrus | 58 | -58 | 28 | 943 | -4.43 | 0.000227 |
| Cuneus | 0 | -76 | 30 | 102 | -4.33 | 0.000256 |
| Middle Cingulum | 4 | -28 | 28 | 77 | -4.29 | 0.000259 |
| Precuneus | 2 | -62 | 38 | 189 | -4.1 | 0.000309 |
| Thalamus | 12 | -24 | 8 | 51 | -4.03 | 0.000309 |
| Calcarine sulcus | -4 | -68 | 22 | 38 | -3.77 | 0.000541 |
| Inderior Temporal gyrus | 52 | -28 | -26 | 26 | -3.76 | 0.000541 |
| Gyrus Rectus | -4 | 38 | -22 | 60 | -3.6 | 0.000689 |
| Thalamus | 4 | -16 | -4 | 97 | -4.53 | 0.000227 |
| Middle Frontal gyrus | 40 | 44 | 36 | 24 | -3.75 | 0.000541 |
| Superior Occipital gyrus | -10 | -96 | 12 | 27 | -3.79 | 0.000541 |
| Middle Temporal gyrus | -52 | -54 | 20 | 48 | -3.88 | 0.000476 |
| Supramarginal gyrus | 52 | -28 | 30 | 20 | -3.25 | 0.001482 |
| Inferior Temporal gyrus | 60 | -38 | -18 | 34 | -3.87 | 0.000476 |
| Middle Occipital gyrus | -38 | -76 | 12 | 21 | -4.01 | 0.000326 |
| Thalamus | -4 | -20 | 6 | 37 | -3.45 | 0.000954 |
| Cuneus | 14 | -78 | 32 | 27 | -3.13 | 0.002079 |
| Fusiform gyrus | -22 | 14 | -48 | 9 | -3.34 | 0.00125 |
| Middle Occipital gyrus | -20 | -76 | 20 | 18 | -3.77 | 0.000541 |
| Postcentral gyrus | 44 | -20 | 32 | 13 | -2.81 | 0.005416 |
| Precuneus | -6 | -72 | 42 | 20 | -3.34 | 0.00125 |
| Inferior Temporal gyrus | 44 | -58 | -8 | 18 | -4.05 | 0.000309 |

| | | | | | | |
|---|---|---|---|---|---|---|
| Middle Cingulum | -2 | -38 | 36 | 14 | -3.46 | 0.000949 |
| Middle Cingulum | 2 | -42 | 32 | 7 | -3.6 | 0.000689 |
| Fusiform gyrus | -18 | 8 | -48 | 7 | -3.22 | 0.001644 |
| Middle Temporal gyrus | -54 | -54 | 6 | 8 | -3.5 | 0.000844 |
| Supramarginal gyrus | 52 | -34 | 40 | 16 | -3.27 | 0.001402 |
| Middle Temporal gyrus | -52 | -48 | -2 | 13 | -3.41 | 0.001052 |
| Parahippocampal gyrus | -12 | 8 | -32 | 2 | -3.09 | 0.002384 |
| Superior Frontal gyrus | -2 | 30 | 50 | 16 | -2.73 | 0.006912 |
| Middle Cingulum | 6 | -44 | 36 | 4 | -3.39 | 0.001104 |
| Middle Cingulum | 6 | -32 | 36 | 5 | -3.51 | 0.000836 |
| Middle Cingulum | 4 | 24 | 40 | 4 | -2.67 | 0.007963 |
| Precuneus | -14 | -52 | 26 | 22 | -3.74 | 0.000541 |
| Inferior Temporal gyrus | -42 | -6 | -50 | 13 | -3.12 | 0.002153 |
| Precuneus | -8 | -62 | 30 | 9 | -3.85 | 0.000491 |

**Supplementary Table 8.** Coordinates of the brain regions showing differences in FC in the dorsal dentate-cortical network in AD>MCI.

| Region | X | Y | Z | Cluster Size | T(182) | pFDR |
|---|---|---|---|---|---|---|
| AD>MCI | | | | | | |
| Middle Frontal Orbital cortex | -48 | 48 | 0 | 24 | 4.24 | 0.000408 |
| Inferior Parietal gyrus | -54 | -60 | 46 | 12 | 4.17 | 0.000408 |
| MCI>AD | | | | | | |
| Middle Temporal gyrus | 68 | -4 | -18 | 29 | -3.66 | 0.001007 |
| Insula cortex | 38 | 22 | 2 | 2 | -3.3 | 0.001649 |
| Precuneus | 20 | -46 | 26 | 7 | -2.81 | 0.006301 |
| Superior Frontal gyrus | -14 | 52 | 20 | 15 | -3.43 | 0.001468 |
| Thalamus | 2 | 0 | 8 | 11 | -3.1 | 0.002736 |
| Angular gyrus | -34 | -52 | 26 | 12 | -2.62 | 0.009633 |
| Superior Temporal gyrus | -64 | -28 | 8 | 3 | -2.98 | 0.003786 |
| Caudate nucleus | -14 | 32 | 6 | 2 | -2.64 | 0.009337 |
| Precentral gyrus | 30 | -12 | 50 | 1 | -2.63 | 0.009599 |
| Postcentral gyrus | 30 | -22 | 50 | 4 | -2.66 | 0.009154 |
| Postcentral gyrus | 30 | -20 | 46 | 1 | -2.72 | 0.007892 |
| Middle Frontal gyrus | 32 | 34 | 26 | 4277 | -4.43 | 0.000345 |
| Middle Frontal gyrus | -20 | 10 | 36 | 1012 | -4.5 | 0.000345 |
| Precuneus | 4 | -54 | 20 | 1483 | -3.84 | 0.000912 |
| Precentral | 34 | -14 | 60 | 1382 | -3.7 | 0.000938 |
| Middle Temporal Pole | 42 | 14 | -32 | 77 | -4.18 | 0.000408 |
| Posterior Cingulum | 0 | -32 | 8 | 277 | -3.51 | 0.001336 |
| Caudate nucleus | -20 | 0 | 26 | 66 | -4.06 | 0.000532 |
| Putamen | 30 | 4 | -4 | 96 | -3.74 | 0.000938 |
| Lingual gyrus | -30 | -62 | -2 | 105 | -3.53 | 0.001336 |

| | | | | | | |
|---|---|---|---|---|---|---|
| Insula cortex | 28 | -14 | 20 | 42 | -3.71 | 0.000938 |
| Superior Temporal gyrus | 38 | -34 | 12 | 109 | -4.02 | 0.000533 |
| Inferior Frontal Operculum | 28 | 10 | 26 | 156 | -3.33 | 0.001551 |
| Angular gyrus | -38 | -48 | 22 | 30 | -3.7 | 0.000938 |
| Middle Temporal gyrus | -52 | -44 | 12 | 54 | -3.48 | 0.001336 |
| Hippocampus | -14 | -34 | 12 | 38 | -3.01 | 0.003539 |
| Thalamus | -10 | -20 | 16 | 23 | -3.22 | 0.00197 |
| Precentral gyrus | 48 | -2 | 50 | 51 | -3.33 | 0.001551 |
| Caudate nucleus | -10 | 18 | 8 | 29 | -3.46 | 0.001391 |
| Middle Cingulum | -14 | -4 | 36 | 30 | -3.54 | 0.001336 |
| Fusiform gyrus | 32 | -40 | -10 | 12 | -3.4 | 0.001468 |
| Hippocampus | 40 | -22 | -8 | 34 | -3.28 | 0.001717 |
| Precentral gyrus | 28 | -16 | 46 | 18 | -3.26 | 0.001758 |
| Superior Frontal gyrus | -10 | 56 | 16 | 4 | -3.39 | 0.001468 |
| Precuneus | -14 | -40 | 8 | 24 | -3.49 | 0.001336 |
| Caudate nucleus | 14 | -2 | 16 | 12 | -3.51 | 0.001336 |
| Putamen | 34 | 4 | -12 | 11 | -3.38 | 0.001468 |
| Hippocampus | 18 | -28 | -4 | 13 | -3.41 | 0.001468 |
| Insula cortex | 34 | 18 | 0 | 8 | -3.36 | 0.00152 |
| Parahippocampal gyrus | 24 | -34 | -8 | 8 | -3.15 | 0.002399 |
| Superior Frontal gyrus | -18 | 48 | 16 | 4 | -3.77 | 0.000938 |

**Supplementary Table 9.** FC connectivity differences in dDN-cortical networks predicting scores for an immediate recall task in AD>CN.

| Region | X | Y | Z | Cluster Size | T(333) | pFDR |
|---|---|---|---|---|---|---|
| AD>CN | | | | | | |
| Superior Temporal gyrus | -50 | 4 | -4 | 14708 | 5.11 | 0.000959 |
| Middle Frontal gyrus | 44 | 46 | 26 | 133 | 4.26 | 0.001737 |
| Superior Temporal gyrus | -62 | -38 | 22 | 146 | 4.15 | 0.001763 |
| Posterior Cingulum | -10 | -30 | 30 | 56 | 3.92 | 0.002072 |
| Inferior Occipital gyrus | -32 | -82 | -2 | 55 | 3.83 | 0.00211 |
| Middle Cingulum | 4 | 0 | 32 | 29 | 3.65 | 0.002326 |
| Putamen | 28 | 10 | 8 | 32 | 3.53 | 0.002389 |
| Precuneus | -10 | -52 | 12 | 16 | 3.4 | 0.002539 |
| Superior Frontal gyrus | 18 | 28 | 32 | 16 | 3.32 | 0.002584 |
| CN>AD | | | | | | |
| Gyrus Rectus | -6 | 38 | -22 | 14543 | -5.21 | 0.000663 |
| Inferior Temporal gyrus | 48 | -44 | -18 | 1459 | -5.07 | 0.000959 |
| Superior Frontal gyrus | -24 | -6 | 60 | 2401 | -4.83 | 0.000959 |
| Cuneus | -2 | -96 | 26 | 1701 | -4.77 | 0.00099 |
| Fusiform gyrus | 24 | -58 | -14 | 357 | -4.71 | 0.00099 |
| Cerebellum Crus I | -30 | -92 | -22 | 435 | -4.43 | 0.001421 |

| | | | | | | |
|---|---|---|---|---|---|---|
| Lingual gyrus | 6 | -86 | -8 | 437 | -4.34 | 0.001441 |
| Cerebellum Crus I | 32 | -86 | -32 | 117 | -4.28 | 0.001709 |
| Supramarginal gyrus | -58 | -24 | 36 | 336 | -4.13 | 0.001774 |
| Hippocampus | 20 | -22 | -4 | 52 | -4.04 | 0.001861 |
| Parahippocampal gyrus | 24 | -16 | -32 | 122 | -4.03 | 0.001917 |
| Thalamus | 4 | -24 | -4 | 55 | -3.96 | 0.001923 |
| Supramarginal gyrus | -46 | -42 | 36 | 185 | -3.96 | 0.001953 |
| Inferior Frontal Orbital cortex | 42 | 24 | -14 | 12 | -3.93 | 0.002032 |
| Posterior Cingulum | -10 | -44 | 32 | 71 | -3.92 | 0.002078 |
| Calcarine sulcus | -18 | -58 | 16 | 50 | -3.75 | 0.002218 |
| Fusiform gyrus | 28 | -82 | -8 | 42 | -3.73 | 0.002239 |
| Paracentral Lobule | -6 | -24 | 70 | 41 | -3.71 | 0.00225 |
| Anterior Cingulum | 10 | 34 | 30 | 10 | -3.71 | 0.002302 |
| Caudate nucleus | 18 | 0 | 20 | 42 | -3.6 | 0.002341 |
| Middle Frontal Orbital cortex | 40 | 62 | -10 | 19 | -3.53 | 0.002389 |
| Calcarine sulcus | -28 | -68 | 8 | 26 | -3.49 | 0.00241 |
| Vermis VI | 2 | -76 | -12 | 15 | -3.45 | 0.00243 |
| Cerebellum Crus I | 36 | -78 | -18 | 31 | -3.44 | 0.002456 |
| Inferior Parietal gyrus | -54 | -52 | 44 | 24 | -3.44 | 0.002461 |
| Middle Frontal gyrus | 26 | 46 | 16 | 9 | -3.43 | 0.002481 |
| Middle Occipital gyrus | -30 | -82 | 16 | 12 | -3.42 | 0.002489 |
| Pars Triangularis of the Inferior Frontal gyrus | 28 | 18 | 26 | 13 | -3.39 | 0.002546 |
| Cerebellum Crus I | -42 | -76 | -18 | 20 | -3.36 | 0.002558 |
| Middle Cingulum | -4 | -26 | 26 | 28 | -3.35 | 0.002575 |
| Vermis X | 4 | -30 | -36 | 7 | -3.31 | 0.002609 |
| Lingual gyrus | -20 | -62 | -4 | 26 | -3.3 | 0.002615 |
| Middle Occipital gyrus | -42 | -72 | 36 | 10 | -3.29 | 0.002627 |
| Caudate nucleus | 16 | 16 | 16 | 33 | -3.23 | 0.002627 |
| Cerebellum III | 10 | -34 | -28 | 5 | -3.23 | 0.002647 |
| Superior Frontal gyrus | 20 | 24 | 36 | 4 | -3.16 | 0.002685 |
| Superior Frontal gyrus | 4 | 62 | 2 | 11 | -3.16 | 0.002685 |
| Anterior Cingulum | 4 | 8 | 30 | 16 | -3.14 | 0.002702 |
| Fusiform gyrus | 44 | -18 | -32 | 4 | -3.11 | 0.002703 |
| Middle Frontal Orbital cortex | 34 | 48 | -18 | 29 | -3.05 | 0.002703 |
| Inferior Occipital gyrus | 42 | -86 | -14 | 6 | -3 | 0.002726 |
| Inferior Temporal gyrus | 48 | -14 | -32 | 9 | -3 | 0.002735 |
| Caudate nucleus | -10 | -14 | 26 | 8 | -2.97 | 0.002735 |
| Inferior Occipital gyrus | 54 | -74 | -14 | 1 | -2.94 | 0.002735 |
| Anterior Cingulum | 14 | 38 | 30 | 2 | -2.88 | 0.002735 |
| Middle Cingulum | -14 | -30 | 40 | 12 | -2.8 | 0.002742 |
| Superior Frontal gyrus | 24 | 52 | 12 | 8 | -2.68 | 0.002759 |
| Superior Frontal gyrus | 10 | 58 | 2 | 8 | -2.34 | 0.002766 |

**Supplementary Table 10.** FC connectivity differences in vDN-cortical networks predicting scores for an immediate recall task in AD>CN.

| Region | X | Y | Z | Size | T(333) | pFDR |
|---|---|---|---|---|---|---|
| AD>CN | | | | | | |
| Supplementary Motor Area | -18 | -2 | 66 | 9576 | 4.98 | 0.003537 |
| Rolandic Operculum | 52 | -6 | 22 | 833 | 4.83 | 0.003537 |
| Cerebellum IV/V | -18 | -40 | -26 | 2146 | 4.7 | 0.003537 |
| Angular gyrus | 42 | -58 | 50 | 218 | 4.61 | 0.003596 |
| Cerebellum IX | -10 | -58 | -52 | 62 | 4.54 | 0.003709 |
| Parahippocampal gyrus | 8 | -6 | -36 | 50 | 4.42 | 0.003839 |
| Middle Frontal gyrus | 28 | 22 | 54 | 20 | 4.07 | 0.003937 |
| Middle Frontal gyrus | 32 | 28 | 50 | 40 | 4.07 | 0.003961 |
| Precuneus | 16 | -52 | 40 | 24 | 4.06 | 0.003988 |
| Inferior Parietal gyrus | 56 | -54 | 40 | 67 | 4.06 | 0.003988 |
| Precuneus | -6 | -54 | 12 | 4 | 3.89 | 0.004059 |
| Inferior Temporal gyrus | -54 | 0 | -36 | 31 | 3.73 | 0.004201 |
| Cerebellum IX | 6 | -40 | -42 | 13 | 3.6 | 0.004238 |
| Middle Occipital gyrus | -44 | -78 | 26 | 16 | 3.52 | 0.004249 |
| Anterior Cingulum | -4 | 28 | 8 | 8 | 3.45 | 0.004276 |
| Supramarginal gyrus | -54 | -44 | 32 | 29 | 3.25 | 0.004276 |
| Hippocampus | 14 | 0 | -12 | 9 | 3.17 | 0.004296 |
| CN>AD | | | | | | |
| Cerebellum IX | -10 | -42 | -56 | 4 | -3 | 0.004296 |
| Calcarine sulcus | -6 | -54 | 8 | 6 | -2.98 | 0.004296 |
| Precuneus | 4 | -48 | 18 | 2 | -2.76 | 0.004296 |
| Parahippocampal gyrus | -6 | -6 | -36 | 8 | -2.75 | 0.004296 |
| Superior Temporal gyrus | 58 | -44 | 20 | 19 | -2.69 | 0.004296 |
| Hippocampus | -28 | -10 | -22 | 1123 | -4.67 | 0.003537 |
| Precuneus | 24 | -48 | 20 | 76 | -4.57 | 0.003709 |
| Middle Cingulum | 0 | -2 | 26 | 49 | -4.52 | 0.003763 |
| Middle Frontal Orbital | -28 | 46 | -10 | 112 | -4.14 | 0.003873 |
| Lingual gyrus | 6 | -64 | 0 | 88 | -3.98 | 0.003988 |
| Supramarginal gyrus | 52 | -44 | 30 | 20 | -3.91 | 0.004059 |
| Vermis X | 0 | -48 | -30 | 44 | -3.89 | 0.004059 |
| Precuneus | 10 | -64 | 42 | 18 | -3.77 | 0.004091 |
| Fusiform gyrus | -34 | -38 | -18 | 18 | -3.77 | 0.004167 |
| Hippocampus | 26 | -6 | -26 | 16 | -3.75 | 0.004201 |
| Amygdala | 32 | 4 | -26 | 19 | -3.7 | 0.004221 |
| Inferior Parietal gyrus | -38 | -38 | 36 | 12 | -3.58 | 0.004238 |
| Cerebellum VIIb | -6 | -76 | -42 | 15 | -3.55 | 0.004238 |
| Heschl gyri | 32 | -32 | 20 | 10 | -3.55 | 0.004238 |
| Superior Frontal gyrus | 20 | 22 | 50 | 9 | -3.52 | 0.004249 |

| | | | | | | |
|---|---|---|---|---|---|---|
| Inferior Parietal gyrus | -52 | -34 | 40 | 27 | -3.49 | 0.004249 |
| Postcentral gyrus | 20 | -38 | 56 | 4 | -3.45 | 0.004276 |
| Precuneus | -14 | -56 | 16 | 10 | -3.45 | 0.004276 |
| Calcarine sulcus | 14 | -62 | 12 | 8 | -3.31 | 0.004276 |
| Precuneus | 4 | -48 | 12 | 6 | -3.23 | 0.004285 |
| Angular gyrus | 34 | -68 | 50 | 4 | -3.2 | 0.004285 |
| Vermis VIII | 0 | -58 | -28 | 4 | -3.19 | 0.004296 |
| Cerebellum IX | 6 | -48 | -46 | 4 | -3.14 | 0.004296 |

**Supplementary Figure 4.** Main Effects of Diagnostic Grouping of both dDN- and vDN-cortical networks. The color bar indicates betas that represent FC. **A.** represents the main effects of dDN-cortical networks. **B.** indicates the main effects of vDN-cortical networks.

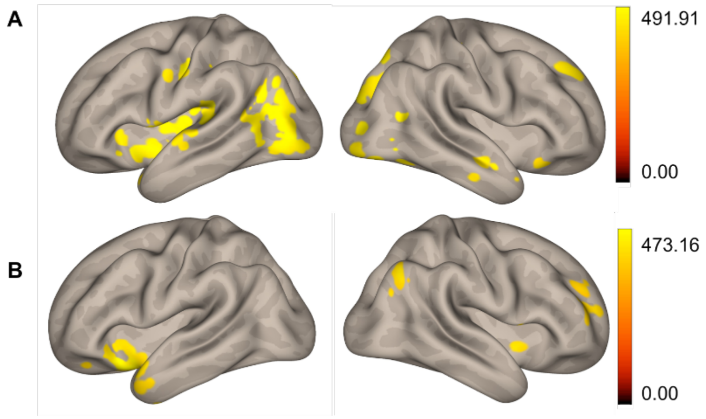


**Supplementary Table 11.** Patterns of differences in FC in AD>MCI associated with an immediate recall task in both dDN- and vDN-cortical networks.

| Region | X | Y | Z | Cluster Size | T(180) | pFDR |
|---|---|---|---|---|---|---|
| **dDN** | | | | | | |
| Cerebellum VIII | -28 | -42 | -58 | 57 | -4.55 | 0.046507 |
| Cerebellum Crus II | -34 | -74 | -48 | 16 | -4.35 | 0.046507 |
| Cerebellum VIII | -28 | -48 | -62 | 17 | -4.26 | 0.046507 |
| Superior Frontal gyrus | -28 | -6 | 60 | 4 | -4.17 | 0.046507 |
| Cerebellum Crus II | -34 | -76 | -44 | 9 | -4.11 | 0.046507 |
| Fusiform gyrus | -38 | -20 | -38 | 8 | -4.11 | 0.046507 |
| Inferior temporal gyrus | -34 | -14 | -48 | 6 | -4.07 | 0.046507 |
| **vDN** | | | | | | |

| | | | | | | |
|---|---|---|---|---|---|---|
| Cerebellum X | -24 | -38 | -38 | 182 | -4.77 | 0.000019 |
| Inferior Temporal gyrus | -38 | 4 | -32 | 252 | 4.32 | 0.000064 |
| Parahippocampal gyrus | -30 | -6 | -28 | 19 | 4.22 | 0.000082 |
| Insula | -34 | 2 | -14 | 28 | 4.09 | 0.000065 |
| Temporal Pole | -42 | 26 | -30 | 5 | 3.55 | 0.000487 |
